# Hypoxia-Mediated Molecular Interactions of Tissue-Specific Mesenchymal Stem Cells Drive Metabolic Reprogramming and Immunomodulation in Acute Graft-versus-Host Disease

**DOI:** 10.1101/2025.01.06.630457

**Authors:** Mohini Mendiratta, Meenakshi Mendiratta, Lakshay Malhotra, Sandeep Rai, Vijaya Sarangathem, Prince Dahiya, Ritu Gupta, Sameer Bakhshi, Vatsla Dadhwal, Deepam Pushpam, Mukul Aggarwal, Aditya Kumar Gupta, Prabhat Singh Malik, Raja Pramanik, Manoranjan Mahapatra, Tulika Seth, Rishi Dhawan, Sabyasachi Bandyopadhyay, GuruRao Hariprasad, Baibaswata Nayak, Thoudam Debraj Singh, Sachin Kumar, Riyaz Ahmed Mir, Surender Kumar Sharawat, Hridayesh Prakash, Sujata Mohanty, Ranjit Kumar Sahoo

## Abstract

**Background:** Mesenchymal stem cells (MSCs) mediate immunomodulation through various mechanisms, including apoptosis, efferocytosis, and mitochondrial transfer. Our study investigates the impact of hypoxia preconditioning on the immune metabolic reprogramming and immunomodulatory potential of MSCs in acute graft-versus-host disease (aGVHD). Additionally, we explored the differential immunomodulatory effects of tissue-specific MSCs, specifically bone marrow (BM) and Wharton’s Jelly (WJ), and elucidated the mechanisms underlying variability in their therapeutic efficacy.

**Methods:** MSCs were isolated from BM and WJ and subjected to hypoxia preconditioning. Their immunometabolic programming potential was assessed by evaluating T-cell proliferation, regulatory T-cell (Treg) induction, effector T-cell differentiation toward Th2, Th9 phenotypes, and macrophage polarization, T-cell bioenergetics in the direct co-culture systems.

**Results:** WJ-MSCs^HYP^ exhibited superior immunomodulatory properties compared to BM-MSCsHYP, by inhibiting T-cell proliferation, enhancing Treg induction, and promoting anti-inflammatory macrophage polarization. WJ-MSCs^HYP^ demonstrated enhanced mitochondrial transfer to T-cell, improving mitochondrial health, reducing ROS, and promoting oxidative phosphorylation, leading to immune homeostasis. Unlike BM-MSCs, WJ-MSCs exhibited higher rates of apoptosis, which facilitated immune modulation through mechanisms independent of efferocytosis.

**Conclusion:** Our findings highlight that WJ-MSCs^HYP^ is a superior candidate for aGVHD by utilizing apoptosis, mitochondrial transfer, and metabolic reprogramming to achieve immune regulation.

## INTRODUCTION

Mesenchymal stem cells (MSCs) have emerged as a pivotal component in immunotherapy due to their remarkable ability to modulate immune responses. These multipotent stromal cells interact dynamically with immune cells within the microenvironment, creating an immunosuppressive milieu conducive to resolving inflammation and tissue repair (1).

Recent studies revealed that MSCs are rapidly cleared from the host system, often through apoptosis induced by the local inflammatory microenvironment in the *in vivo* milieu (2). Cytotoxic T lymphocytes (CTLs), actively induce their apoptosis (3) and this apoptotic process is a byproduct of their limited survival and a critical feature that drives their immunomodulatory efficacy. Apoptotic MSCs are engulfed by phagocytic cells, such as macrophages and dendritic cells, through a process known as efferocytosis. This interaction reprograms these immune cells towards a more anti-inflammatory or tolerogenic phenotype and supports immune homeostasis (4). Unlike viable MSCs, which may evoke variable immune responses depending on the host’s immune profile (5,6), apoptotic MSCs consistently create an immunosuppressive environment, making them particularly effective in conditions such as graft-versus-host disease (GVHD).

While bone marrow (BM) remains the gold standard source for MSCs and is the focus of most preclinical and clinical studies, our findings suggest that Wharton’s Jelly-derived MSCs (WJ-MSCs) exhibit superior immunomodulatory efficiency (4,7). This enhanced efficacy may be attributed to their unique propensity for apoptosis and efficient clearance through efferocytosis, setting them apart from BM-MSCs in their ability to modulate immune responses.

To enhance the immunomodulatory potential and overcome variability, hypoxia preconditioning has emerged as a critical strategy. Hypoxic environments mimic the physiological conditions of inflamed tissues, activating hypoxia-inducible factors (HIFs) in MSCs, which in turn upregulate the secretion of anti-inflammatory cytokines, extracellular vesicles, and apoptotic signals (4). Additionally, hypoxia preconditioning might prime MSCs for efficient apoptosis and efferocytosis, enhancing their capacity to modulate immune cells toward an anti-inflammatory phenotype.

MSCs play a pivotal role in modulating the mitochondrial health and metabolic state of immune cells through mitochondria transfer (8), thereby alleviating oxidative stress in inflammatory conditions. Beyond mitigating oxidative stress, MSCs-derived mitochondria induce metabolic reprogramming of immune cells, thereby, maintaining immune homeostasis (9,10). This dual effect—restoring mitochondrial health and reprogramming immune cell metabolism highlights the multifaceted immunomodulatory potential of MSCs, extending beyond cytokine secretion to directly address mitochondrial dysfunction in inflamed environments.

Given this, it becomes essential to explore the variability in immune responses mediated by MSCs based on their tissue of origin, with particular emphasis on how apoptosis and efferocytosis contribute to differences in immune regulation. Furthermore, understanding how hypoxia preconditioning enhances the immunomodulatory capacity of MSCs by modulating metabolic reprogramming through mitochondrial transfer is crucial for improving their therapeutic immunomodulatory effectiveness in GVHD.

To the best of our knowledge, this is the first study to investigate the variability in immune responses by directly examining and correlating it with the ability of MSCs (naïve, hypoxia-preconditioned; BM, WJ) to undergo apoptosis and efferocytosis under *in vitro* conditions. Moreover, we comprehensively understand how these processes influence immune regulation. This novel approach highlights the critical role of apoptosis and efferocytosis in shaping the immunomodulatory potential of MSCs, offering new insights into the variability of their effects in different tissue sources. Additionally, we have investigated the role of mitochondrial transfer in immune metabolic reprogramming, demonstrating how it contributes to the establishment of a robust immunosuppressive environment in GVHD.

## MATERIAL AND METHODS

### Ethical approval

The study involved human subjects and received approval from the Institutional Human Ethics Committee at the All India Institute of Medical Sciences, New Delhi, India (Ref. No. IECPG-542/23.09.2020). Additionally, it included the use of human MSCs, which were approved by the Institutional Committee for Stem Cell Research at the same institution [Ref. No.: IC-SCR/110/20(R)].

Informed written consent was obtained from the participants, and all procedures followed the guidelines and regulations approved by the ethics committee.

### Isolation, characterization, and hypoxia-preconditioning of MSCs from human bone marrow (BM) and Wharton’s Jelly (WJ)

Human BM aspirates were obtained from ten healthy donors who were recipients of allogeneic stem cell transplants at the Department of Medical Oncology, Dr. B. R. Ambedkar Institute Rotary Cancer Hospital, AIIMS, New Delhi. Additionally, human umbilical cord (UC) tissue was collected from ten donors in sterile transport media containing 1000 IU/ml heparin (Gland Pharma Limited, India) and 200 μg/ml gentamicin (Thermo Fisher Scientific, USA) from the Department of Obstetrics and Gynaecology at AIIMS, New Delhi. BM-MSCs and WJ-MSCs were isolated following our previously established protocol, as described elsewhere (Mendiratta et al., 2024). Both BM-MSCs and WJ-MSCs (Passage-3) were characterized for their plastic adherence, surface marker profile, and trilineage differentiation potential according to the International Society for Cellular Therapy (ISCT) guidelines (11) using our established protocols (4). Passages 3-5 were used for subsequent *in vitro* experiments and these were pooled together at their respective passages.

MSCs were preconditioned with 1%O_2_ in 1X LG-DMEM complete media for 24 hours in a tri-gas incubator (Thermo Fisher Scientific, USA), termed MSCs^HYP^ (4).

### Karyotyping of MSCs

The cells were first expanded in culture to a sufficient number, typically 2*10^6^ cells/ml, in 1X LG-DMEM complete medium (Thermo Fisher Scientific, USA) at 37°C, 5% CO₂. To synchronize the cells in metaphase, they were treated with colchicine (0.1 µg/ml) (Thermo Fisher Scientific, USA) for 6 hours. After colchicine treatment, the cells were harvested using 0.05% Trypsin-EDTA (Thermo Fisher Scientific, USA), centrifuged at 800 rpm for 5 minutes, and resuspended in a hypotonic solution of 0.075 M KCl (Sisco Research Laboratories Pvt. Ltd., India) for 30 minutes at 37°C. After centrifugation, the cells were fixed by adding fixative [methanol (Sisco Research Laboratories Pvt. Ltd., India): acetic acid (Sisco Research Laboratories Pvt. Ltd., India) in a 3:1 ratio] dropwise, incubated for 10 minutes, and this process was repeated 2-3 times. Microscope slides were prepared by dropping the fixed cell suspension onto clean slides and allowing them to air dry. The slides were stained with Giemsa solution (Sisco Research Laboratories Pvt. Ltd., India) at 1:5 dilution for 15 minutes, rinsed with distilled water, and air dried. The stained slides were observed under a light microscope (Zeiss Axio Imager 2, Zeiss, Germany) at 100x magnification to visualize the metaphase chromosomes. The slides were scanned, images were captured using Metafer software (MetaSystems, Germany), and the resulting karyotype was generated using IKAROS software (MetaSystems, Germany) (12).

### *In vitro* T-cell proliferation assay

Peripheral blood (PB) was collected from grade II-IV aGvHD patients (n=25) in sterile sodium heparin-coated vacutainers (BD Biosciences, US). The isolation of peripheral blood mononuclear cells (PBMNCs) was performed through Ficoll density gradient centrifugation (María Del Pilar De la Rosa-Ruiz et al, 2019) and subsequently, CD3^+^ T-cell were extracted from the PBMNCs through negative selection using a Pan T cell isolation kit (Miltenyi Biotec, USA), stained with 1 µM cell Trace^TM^ carboxyfluorescein succinimidyl ester (CFSE) dye (BD Biosciences, USA) and activated with PHA (1µg/ml) (Sigma, USA) and IL-2 (50IU/ml) (Thermo Fisher Scientific, USA) for 48 hours according to our established protocols (4).

For co-culture experiments, tissue-specific (BM, WJ) MSCs (MSCs, MSCs^HYP^) were treated with mitomycin-C at a concentration of 15µg/ml (Thermo Fisher Scientific, USA) for 1 hour in 1X LG-DMEM incomplete medium. Following this treatment, the mitomycin-treated MSCs were co-cultured with CFSE-labeled activated T-cell at a ratio of 1:10 in a mixture of 1X RPMI-1640 complete medium and 1X LG-DMEM complete medium at a 1:1 ratio for 3 days. After the co-culture period, T cell proliferation was evaluated using a DxFlex flow cytometer (Beckman Coulter, USA), and the data analysis was performed using Kaluza software version 2.1 (Beckman Coulter, USA). The normalization of T-cell proliferation in the direct co-culture was performed with CFSE-labeled activated T-cell only (4).

### Induction of regulatory T-cell (Tregs)

The mitomycin-treated MSCs and activated T-cell were co-cultured at a ratio of 1:10 for 5 days. Following this period, the cells were collected, washed with 1X PBS, and stained with fluorochrome-conjugated anti-human monoclonal antibodies targeting CD3, CD4, CD8, CD25, and CD127 (Beckman Coulter, USA). A minimum of 50,000 events were acquired using a DxFlex flow cytometer (Beckman Coulter), and the data were analyzed with Kaluza software version 2.1 (Beckman Coulter). Activated T-cell cultured without MSCs served as a control to establish baseline expression levels of CD3^+^ CD4^+^ CD25^+^ FOXP3^+^ Tregs (4,13).

### Enumeration of effector memory helper T cell subtypes (Th1, Th2, Th9, and Th17)

The proportions of Th1, Th2, Th9, and Th17 cells were quantified in the 5-day co-culture of mitomycin-treated MSCs and activated T-cell by staining with fluorochrome-conjugated anti-human monoclonal antibodies specific for CXCR3, CXCR5, CCR10, CCR4, CCR7, and CCR6 (BD Biosciences, USA) at 37°C for 30 minutes. This was followed by additional staining with anti-human fluorochrome-conjugated CD3, CD4, CD8, and CD45RA monoclonal antibodies. A minimum of 50,000 cells were acquired using a DxFlex flow cytometer (Beckman Coulter, USA) and the data was analyzed using Kaluza software version 2.1 (Beckman Coulter, USA). Activated T-cell cultured without MSCs served as a control to establish baseline expression levels of Th1, Th2, and Th17 (4).

### Enumeration of αβ and γδ CD4^+^ T-cell

The proportions of αβ and γδ CD4^+^ T-cell was quantified in the 3-day co-culture of mitomycin-treated MSCs and activated T-cell by staining with fluorochrome-conjugated anti-human monoclonal antibodies specific for CD3, CD4, CD8, αβ and γδ monoclonal antibodies for 30 minutes in dark. A minimum of 50,000 cells were acquired using a DxFlex flow cytometer (Beckman Coulter, USA) and the data was analyzed using Kaluza software version 2.1 (Beckman Coulter, USA). Activated T-cell cultured without MSCs served as a control to establish baseline expression levels of αβ and γδ T-cell (14).

### Macrophage polarization

CD14^+^ monocytes were isolated from PBMNCs using a pan-monocyte isolation kit (Miltenyi Biotec, USA), as described elsewhere (Mendiratta et al, 2024). Mitomycin-treated MSCs were then co-cultured with the M1 macrophages at a ratio of 1:10 for 3 days. The polarization of macrophages from the M1 to M2 phenotype was assessed by staining the cells with fluorochrome-conjugated antibodies against CD206 (BD Biosciences, USA) for 30 minutes. Following surface staining, cells were fixed and permeabilized using an IntraPrep permeabilization kit (Beckmann Coulter, USA) followed by intracellular staining of cells with fluorochrome-conjugated anti-human iNOS, Arginase-I (Thermo Fisher Scientific, USA) for 30 minutes. The cells were acquired using a DxFlex flow cytometer (Beckman Coulter, USA) to enumerate the M1 and M2 macrophages (4).

### Estimation of secreted cytokines, chemokines, and other soluble factors

The concentration of various secreted factors, including, Prostaglandin E2 (PGE2), Interleukin-10 (IL-10), Interferon-γ (IFN-γ), Tumor necrosis factor-α (TNF-α), Interleukin-12 (IL-12), Transforming growth factor-β (TGF-β), Interleukin-6 (IL-6), and Interleukin-1β (IL-1β) (Thermo Fisher Scientific, USA) were quantified in the culture-conditioned media (CCM) of co-culture of mitomycin-treated MSCs and aPBMNCs using ELISA with a PR4100 microplate reader (Bio-Rad, USA) following manufacturer’s instructions. CCM from the culture of aPBMNCs alone (without co-culture) served as a control for baseline expression of secreted soluble factors (15).

### Kyneurine assay

The kynurenine assay evaluated the IDO activity in the direct co-culture of MSCs (MSCs, MSCs^HYP^) and aPBMNCs. Following 3 days of co-culture, 200 µl of the culture supernatant was collected from each well and deproteinized by adding an equal volume of 30% trichloroacetic acid (TCA) (Sigma-Aldrich, Saint Louis, MO, USA). The mixture was vortexed, and centrifuged at 10,000 × g for 10 minutes at 4°C, and the clear supernatant was transferred to a new Eppendorf tube (Genaxy Scientific, India). For kynurenine detection, 100 µl of the deproteinized supernatant was mixed with an equal volume of Ehrlich’s reagent (Sigma-Aldrich, Saint Louis, MO, USA) in a 96-well plate (Genaxy Scientific, India) and incubated at room temperature for 10 minutes to develop a yellow color. The absorbance was measured at 490 nm using a PR4100 microplate reader (Bio-Rad, USA), and kynurenine levels were quantified by interpolating the absorbance values from a standard curve prepared using known concentrations of kynurenine. CCM from the culture of aPBMNCs alone (without co-culture) served as a control for the baseline expression of secreted IDO (16).

### Annexin-V/7AAD assay

The proportion of apoptosis of MSCs was assessed in the 3-day co-culture of mitomycin-treated MSCs and activated T-cell using Annexin-V/7AAD (BD Biosciences, USA) staining to assess the effect of T-cell on MSCs (4).

### *In vitro* MSCs phagocytosis assay

CFSE-labeled aPBMNCs were treated with pH Rhodo red succinimidyl ester-labeled MSCs (Thermo Fisher Scientific, USA) at a 1:10 ratio for 72 hours. The percentage of CD14^+^ monocytes that were CFSE-positive also exhibited the pH Rhodo red signal, which was utilized to calculate the engulfed MSCs by the monocytes using a DxFlex flow cytometer (Beckman Coulter, USA) (4).

### T-cell phenotype assay

The subtypes of CD4^+^ and CD8^+^ T-cell were assessed in the 3-day co-culture of mitomycin-treated MSCs and aPBMNCs by staining the cell suspension with fluorochrome-conjugated anti-human CCR7 (Beckmann Coulter, USA) monoclonal antibodies at 37֯C for 30 minutes, followed by surface staining with anti-human CD3, CD4, CD8, and CD45RA (Beckmann Coulter, USA) monoclonal antibodies for 30 minutes at room temperature. A minimum of 50,000 cells were acquired using a DxFlex flow cytometer (Beckman Coulter, USA) and the data was analyzed using Kaluza software version 2.1 (Beckman Coulter, USA). Activated T-cell cultured without MSCs served as a control to establish baseline expression levels of subtypes of CD8^+^ T-cell (17).

### Mitochondria transfer assay

The transfer of mitochondria from MSCs to CD3^+^ T-cell was assessed in the direct co-culture of mitomycin-treated MSCs and aPBMNCs. Briefly, MSCs were pre-labeled with Mitotracker^TM^ Red (Thermo Fisher Scientific, USA) at a concentration of 100nM, and aPBMNCs were pre-labeled with Cell Tracker^TM^ Green (Thermo Fisher Scientific, USA) at a concentration of 1μM at 37֯C for 20 minutes followed by their direct co-culture for 24 hours. The proportion of Mitotracker^+^ Cell Tracker Green^+^ cells was enumerated to assess the transfer of mitochondria from MSCs to CD3^+^ T-cell using a DxFlex flow cytometer (Beckmann Coulter, USA) (18).

### Quantification of mitochondrial DNA (mtDNA) copy number

Genomic and mitochondrial DNA were isolated from MSCs and MSCs^HYP^ before co-culture, as well as from T-cell before and after co-culture for the assessment of mtDNA copy number using a QIAamp DNA Mini Kit (Qiagen, Netherlands), according to the manufacturer’s instructions. Real-time PCR was performed with 50 ng of DNA, 5 μl of Kappa SYBR Master Mix, and primers at a final concentration of 0.3 μM in a total reaction volume of 10 μl using a CFX96 Real-Time System (Bio-Rad). The copy number of mtDNA and nuclear DNA was quantified using the threshold cycle (Ct) values. The Delta Ct (ΔCt) was determined using the equation: Ct (mitochondrial gene) − Ct (nuclear gene) and the relative mtDNA copy number was calculated using the 2−ΔΔCt method (18).

### Measurement of mitochondrial membrane potential

The mitochondrial health of CD3^+^ T-cell was assessed in the 1-day co-culture of mitomycin-treated MSCs and CD3^+^ T-cell by staining the cell suspension with JC-1 dye (Thermo Fisher Scientific, USA) at a concentration of 2μM at 37֯C for 20 minutes in the dark. Subsequently, the cells were washed with 1XPBS and stained with fluorochrome-conjugated anti-human CD3 monoclonal antibody. JC-1 aggregates/JC-1 monomers ratio gated on CD3^+^ T-cell was enumerated to assess their mitochondrial health status (19).

### Mitochondrial Reactive Oxygen Species (ROS) assay

The mitochondrial ROS level of CD3^+^ T-cell was assessed in the 1-day co-culture of CD3^+^ T-cell and mitomycin-treated MSCs by staining the cell suspension with MitoSOX Red (Thermo Fisher Scientific, USA) at a concentration of 5μM at 37֯C for 20 minutes. Subsequently, the cells were washed with 1XPBS and stained with fluorochrome-conjugated anti-human CD3 monoclonal antibody. The percentage of CD3^+^ MitoSox Red^+^ cells was quantified to enumerate the mitochondrial ROS level in CD3^+^ T-cell using a DxFlex flow cytometer (Beckmann Coulter, USA), and the data was analyzed using Kaluza Software Version 2.1. CD3^+^ T-cell cultured without MSCs served as a control to establish baseline expression of mitochondrial ROS (20).

### T-cell mitochondrial bioenergetics

Oxygen consumption rate (OCR) and extracellular acidification rate (ECAR) were assessed using the Seahorse XFe24 Extracellular Flux Analyzer (Agilent Technologies, USA), which serves as indicators for oxidative phosphorylation (OXPHOS) and glycolysis, respectively. Mitomycin-treated MSCs were co-cultured with CD3^+^ T-cell for 24 hours followed by the separation of CD3^+^ T-cell from the cell suspension of co-culture using negative selection.

Further, these co-cultured CD3^+^ T-cell and control (non-co-cultured) CD3^+^ T-cell were seeded separately at a density of 1*10^6^ cells per well on CellTak-coated plates (Corning) in XF-Base Media (Agilent Technologies, USA), supplemented with 2.5 mM glucose, 1 mM sodium pyruvate (HiMedia, India), and 2 mM L-glutamine (HiMedia, India). Control CD3^+^ T-cell was used as a baseline for assessing T-cell mitochondrial bioenergetics. The OCR measurements were taken sequentially, starting from the basal level, followed by the addition of 1.0 mM oligomycin (Sigma-Aldrich, Saint Louis, MO, USA), 0.75 mM FCCP (fluorocarbonyl cyanide phenylhydrazone) (G-Biosciences, Saint Louis, MO, USA), and a combination of 0.5 mM rotenone (Sigma-Aldrich, Saint Louis, MO, USA) and antimycin (Sigma-Aldrich, Saint Louis, MO, USA) to evaluate changes in mitochondrial respiratory parameters. Data analysis was performed using Wave 2.6.1 and GraphPad Prism software (21).

### Identification of secreted proteins using liquid chromatography-assisted mass spectrometry (LC-MS/MS)

Protein per sample (BM-MSCs, WJ-MSCs, and their co-culture with aPBMNCs) was used for digestion and reduced with 5 mM tris(2-carboxyethyl) phosphine (TCEP) and further alkylated with 50 mM iodoacetamide and then digested with Trypsin (1:50, Trypsin/lysate ratio) for 16 h at 37 °C. Digests were cleaned using a C18 silica cartridge to remove the salt and dried using a speed vac. The dried pellet was resuspended in buffer A (2% acetonitrile, 0.1% formic acid).

Experiments were performed on an Easy-nlc-1000 system coupled with an Orbitrap Exploris mass spectrometer. 1µg of peptide sample was loaded on C18 column 15 cm, 3.0μm Acclaim PepMap (Thermo Fisher Scientific, USA) and separated with a 0– 40% gradient of buffer B (80% acetonitrile, 0.1% formic acid) at a flow rate of 500 nl/min) and injected for MS analysis. LC gradients were run for 110 minutes. MS1 spectra were acquired in the Orbitrap (Max IT = 60ms, AGQ target = 300%; RF Lens = 70%; R=60K, mass range: 375−1500; Profile data). Dynamic exclusion was employed for 30s excluding all charge states for a given precursor. MS2 spectra were collected for the top 20 peptides. MS2 (Max IT= 60ms, R= 15K, AGC target 100%).

All samples were processed and RAW files generated were analyzed with Proteome Discoverer (v2.5) against the UniProt Human database. For dual Sequest and Amanda search, the precursor and fragment mass tolerances were set at 10 ppm and 0.02 Da, respectively. The protease used to generate peptides, i.e. enzyme specificity was set for trypsin/P (cleavage at the C terminus of “K/R: unless followed by “P”). Carbamidomethyl on cysteine as fixed modification and oxidation of methionine and N-terminal acetylation were considered as variable modifications for database search. Both peptide spectrum match and protein false discovery rate were set to 0.01 FDR (22)

### Statistical analysis

All statistical analyses were conducted using GraphPad Prism version 8.4.3. One-way and Tukey’s post hoc tests compared three or more groups. Data was shown as Mean±S.D. and a p-value of ≤ 0.05 was considered statistically significant.

## RESULTS

### Hypoxia preconditioning of MSCs preserved their parental identity

Human MSCs isolated from both BM and WJ retained their parental identity following hypoxic preconditioning (1% O₂, 24 hours). These cells maintained their characteristic spindle-shaped fibroblast-like morphology (Figure S1A) and expressed ≥95% of the surface markers CD105, CD73, CD90, CD29, and HLA-I, with minimal or no expression (≤2%) of lineage-specific markers such as CD34 and CD45, as well as HLA-II (Figure S1B, C). Additionally, both MSCs populations demonstrated their *in vitro* potential to differentiate into mesodermal lineages (Figure S1D). Thus, 1% O_2_ did not affect the core properties of MSCs, regardless of their tissue origin.

MSCs exhibited a normal karyotype, as confirmed by chromosomal analysis. Notably, BM-MSCs displayed a chromosomal complement of 44 autosomes along with XY sex chromosomes, consistent with a normal male karyotype and WJ-MSCs exhibited 44 autosomes with XX sex chromosomes, indicative of a normal female karyotype (Figure S1E).

### WJ-MSCs^HYP^ inhibited CD3^+^ T-cell proliferation, polarized effector CD4^+^ T-cell to a regulatory and anti-inflammatory phenotype

The immunomodulatory effects of MSCs derived from BM and WJ, both in their naïve (MSCs) and hypoxia-preconditioned (MSCs^HYP^) states, were evaluated in their direct co-culture with aGVHD patients derived CD3^+^ T-cell to assess their impact on T-cell proliferation and polarization towards anti-inflammatory or regulatory phenotype (Figure 1A).

**Figure 1:**
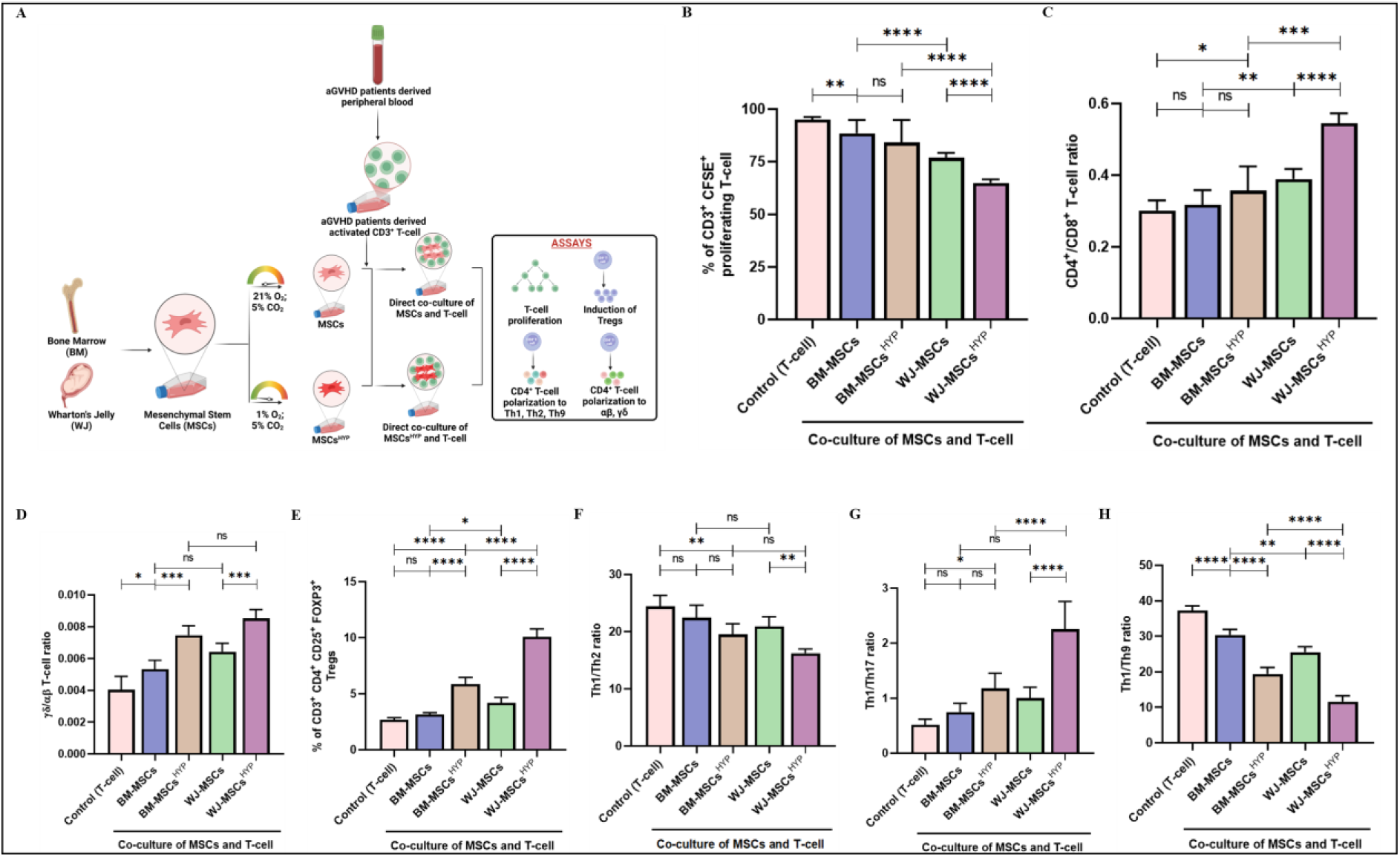
Effect of naïve and hypoxia-preconditioned MSCs (MSCs, MSCs^HYP^) on the aGVHD patients derived T-cell reprogramming. (A) Schematic representation of the co-culture model for the assessment of T cell programming. The bar graph represents (B) the percentage of CD3^+^ CFSE^+^ proliferating T-cell (n=25). (C) the ratio of CD4^+^/CD8^+^ T-cell (n=25). (D) the ratio of γδ/αβ CD4^+^ T-cell (n=25). (E) the percentage of CD3^+^ CD4^+^ CD25^+^ FOXP3^+^ Tregs (n=25). (F) the ratio of Th1/Th2 (n=25). (G) the ratio of Th1/Th17 (n=25). (H) the ratio of Th1/Th9 (n=25) in the direct co-culture of MSCs and T-cell. Data shown represent the Mean±S.D of 25 independent experiments performed with T-cell derived from 25 different donors (biological replicates), with each experiment conducted in triplicate (technical replicates). Statistical analysis: Tukey’s multiple comparisons test; *≤0.05; **≤0.01; ***≤0.001; ****≤0.0001. Abbreviations: BM: Bone marrow; WJ: Wharton’s Jelly; MSCs: Mesenchymal Stem Cells; HYP: Hypoxia-preconditioned

We observed that both MSCs inhibited the proliferation of CD3^+^T-cell, with WJ-MSCs demonstrating greater efficacy than BM-MSCs (88.552% vs. 76.851%; p: ≤0.0001). Hypoxia preconditioning enhanced their immunosuppressive capabilities, regardless of tissue origin. However, no significant differences were observed between BM-MSCs and BM-MSCs^HYP^ (p: 0.1153) while WJ-MSCs^HYP^ exhibited a more profound effect than WJ-MSCs (64.956% vs. 76.851%; p: ≤0.0001). Interestingly, WJ-MSCs^HYP^ was superior to BM-MSCs^HYP^ (65.62% vs. 84.134%; p: ≤0.0001) in suppressing CD3^+^ T-cell proliferation (Figure 1B).

There was no significant difference was observed in the CD4^+^/CD8^+^ T-cell ratio between control and BM-MSCs (p: 0.914) and BM-MSCs and BM-MSCs^HYP^ (p: 0.239). However, WJ-MSCs demonstrated a significantly higher CD4+/CD8+ T-cell ratio compared to BM-MSCs (0.389 vs. 0.3176; p: ≤0.01). Similarly, WJ-MSCs^HYP^ exhibited superior efficacy in enhancing the CD4^+^/CD8^+^ T-cell ratio compared to BM-MSCs^HYP^ (0.545 vs. 0.357; p: ≤ 0.001). These findings suggest that MSCs, particularly WJ-MSCs and WJ-MSCs^HYP^, selectively suppress CD8^+^ T-cell proliferation, leading to a skewed T-cell response (Figure 1C).

In the direct co-culture system, MSCs exhibited a distinct ability to modulate the proportions of αβ and γδ T-cell subsets within the CD4^+^ T-cell population. Specifically, MSCs reduced the proportion of αβ CD4^+^ T-cell while increasing the γδ subset. BM-MSCs^HYP^ demonstrated significantly greater efficacy compared to their naïve counterparts (0.0075 vs. 0.0054; p: ≤ 0.0001), and a similar trend was observed with WJ-MSCs^HYP^ versus WJ-MSCs (0.0085 vs. 0.0064; p: ≤ 0.0001). These findings highlight the enhanced immunomodulatory potential of MSCs through hypoxia preconditioning, irrespective of their tissue origin. While WJ-MSCs^HYP^ appeared superior to BM-MSCs^HYP^ in increasing the γδ proportion (0.0085 vs. 0.0075) (Figure 1D), the difference was not statistically significant (p: 0.098), suggesting that while WJ-MSCs^HYP^ may offer incremental advantages, both sources are comparably effective when preconditioned under hypoxia.

These findings raised the question of whether MSCs promote the polarization of CD4^+^ T-cell toward a regulatory or anti-inflammatory phenotype, highlighting the need to delineate their specific effects on CD4^+^ T-cell subsets. Our results showed that MSCs induced Tregs generation, with no significant increase in Tregs induction by BM-MSCs compared to control (p: 0.576). In contrast, BM-MSCs^HYP^ exhibited a significantly greater effect on Treg generation compared to BM-MSCs (5.86% vs. 3.15%; p: ≤0.0001). Notably, WJ-MSCs^HYP^ showed markedly enhanced capacity to induce Tregs compared to WJ-MSCs (10.08% vs 4.21%; p: ≤0.0001) (Figure 1E, S4A-B). Furthermore, our findings demonstrated that MSCs promoted the polarization of CD4^+^ T-cell toward an anti-inflammatory Th2 phenotype, as indicated by a reduced Th1/Th2 ratio. The Th1/Th2 ratio showed no significant difference, neither between BM-MSCs and the control group (22.42 vs 24.42; p: 0.4098) nor between BM-MSCs and BM-MSCs^HYP^ (22.424 vs 19.52; p: 0.1089). In contrast, WJ-MSCs^HYP^ exerted a stronger effect in reducing the Th1/Th2 ratio compared to WJ-MSCs (16.25 vs 20.88; p: ≤0.01) (Figure 1F). A similar trend was observed with the Th1/Th17 ratio, wherein WJ-MSCs^HYP^ demonstrated significantly a greater decrease in Th17 compared to BM-MSCs^HYP^ (2.252 vs 1.178; p: ≤0.0001) (Figure 1G). Both MSCs and MSCs^HYP^ demonstrated a notable ability to polarize CD4^+^ T-cell toward the anti-inflammatory Th9 phenotype. WJ-MSCs were significantly more effective than BM-MSCs (25.526 vs 30.338; p: ≤0.01) and their hypoxia preconditioned counterparts (WJ-MSCs^HYP^) also outperformed BM-MSCs^HYP^ (11.598 vs 19.42; p: ≤0.0001) (Figure 1H).

These findings suggested that hypoxia preconditioning improved the functionality of MSCs, regardless of their tissue source, though WJ-MSCs may have intrinsic advantages. This makes them a promising candidate for therapeutic strategies aiming to balance immune suppression and tumor surveillance in transplant recipients.

### WJ-MSCs^HYP^ promoted macrophage polarization toward M2 phenotype

Certainly, T-cell are key players in GVHD; however, antigen-presenting cells (dendritic cells and macrophages) also contribute to GVHD progression through their interactions with T-cell, which amplify the cytokine storm (23). To study the impact of naïve or MSC^HYP^ on aGVHD patients-derived antigen-presenting cells (Figure 2A), we assessed macrophage polarization in a direct co-culture system using flow cytometry. Our results revealed that both MSCs and MSCs^HYP^ exhibited a pronounced effect on macrophage polarization under *in vitro* conditions. We did not observe any significant difference in the inhibition of the M1 phenotype between BM-MSCs^HYP^ and BM-MSCs (78.027% vs 83.564%; p: 0.250). However, a significant difference was observed between WJ-MSCs and WJ-MSCs^HYP^ (66.958% vs 57.639%; p: ≤0.01). Interestingly, both WJ-MSCs and WJ-MSCs^HYP^ significantly outperformed BM-MSCs and BM-MSCs^HYP^ counterparts, respectively, in inhibiting M1 phenotype (66.96% vs 83.56%; p: ≤0.0001; 57.64% vs 78.03%; p: ≤0.0001) (Figure 2B).

**Figure 2:**
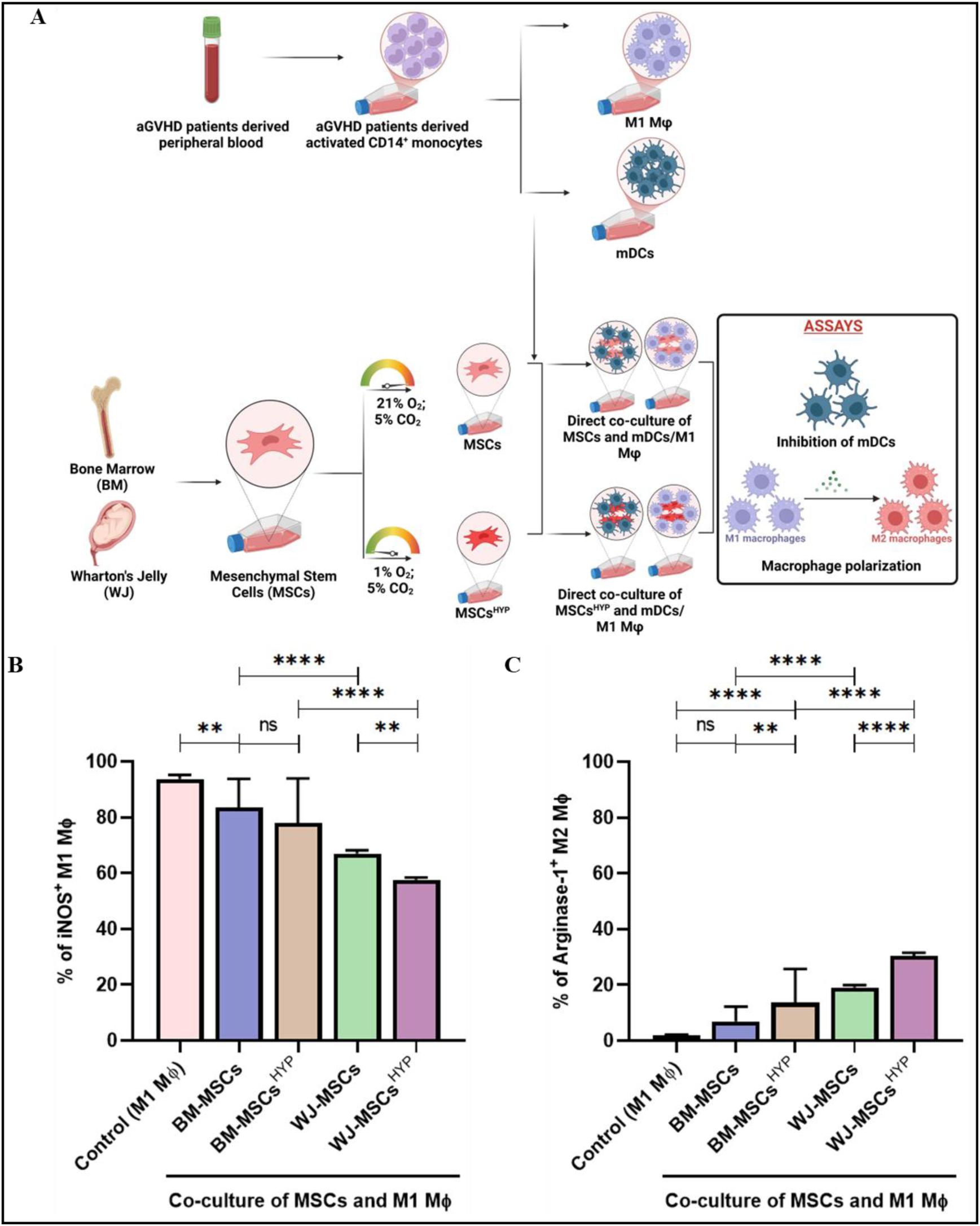
Effect of naïve and hypoxia-preconditioned MSCs (MSCs, MSCs^HYP^) on the antigen-presenting cells of aGVHD patients. (A) Diagrammatic representation of the co-culture model for the evaluation of impact of MSCs/MSCs^HYP^ on antigen-presenting cells. The bar graph represents (B) the percentage of iNOS^+^ M1 MФ. (C) the percentage of Arginase-1^+^ M2 MФ in the direct co-culture of MSCs and M1 MФ. Data shown represent the Mean±S.D of 25 independent experiments performed with mDCs/ M1 MФ derived from 25 different donors (biological replicates), with each experiment conducted in triplicate (technical replicates). Statistical analysis: Tukey’s multiple comparisons test; *≤0.05; **≤0.01; ***≤0.001; ****≤0.0001. *Abbreviations: BM: Bone marrow; WJ: Wharton’s Jelly; MSCs: Mesenchymal Stem Cells; HYP: Hypoxia-preconditioned; mDCs: Mature Dendritic Cells*

A similar trend was observed in polarizing macrophages toward an M2 anti-inflammatory phenotype, with WJ-MSCs^HYP^ exhibiting superior efficacy compared to BM-MSCs^HYP^, as evidenced by a more pronounced increase in the proportion of Arginase-1⁺ M2 macrophages (30.491% vs. 13.662%; p ≤ 0.0001) (Figure 2C).

### WJ-MSCs^HYP^ enhanced IDO, PGE2, IL-10, and TGF-β secretion while inhibiting IFN-γ, TNF-α, IL-12, IL-6 and IL-1β

Our experiments demonstrated that WJ-MSCs^HYP^ demonstrated a superior immunomodulatory profile compared to BM-MSCs^HYP^. Specifically, WJ-MSCs^HYP^ exhibited significantly enhanced secretion of key immunoregulatory factors, including IDO, PGE2, IL-10, and TGF-β. These molecules are critical mediators of anti-inflammatory responses, suggesting a heightened potential of WJ-MSCs^HYP^ to modulate immune responses favorably. Conversely, WJ-MSCs^HYP^ showed greater inhibition of pro-inflammatory cytokines and chemokines, including IFN-γ, TNF-α, IL-6, and CCR6, compared to BM-MSCs^HYP^. These findings indicate that WJ-MSCs^HYP^ are more effective at suppressing inflammatory pathways, particularly those mediated by Th1 and Th17 cells, which are key drivers of immune dysregulation in aGVHD. Although hypoxic preconditioning enhanced the immunomodulatory capabilities of both BM-MSCs and WJ-MSCs compared to their non-hypoxic counterparts, BM-MSCs^HYP^ was not as pronounced as the comparative superiority observed over WJ-MSCs (Figure 3A-H).

**Figure 3:**
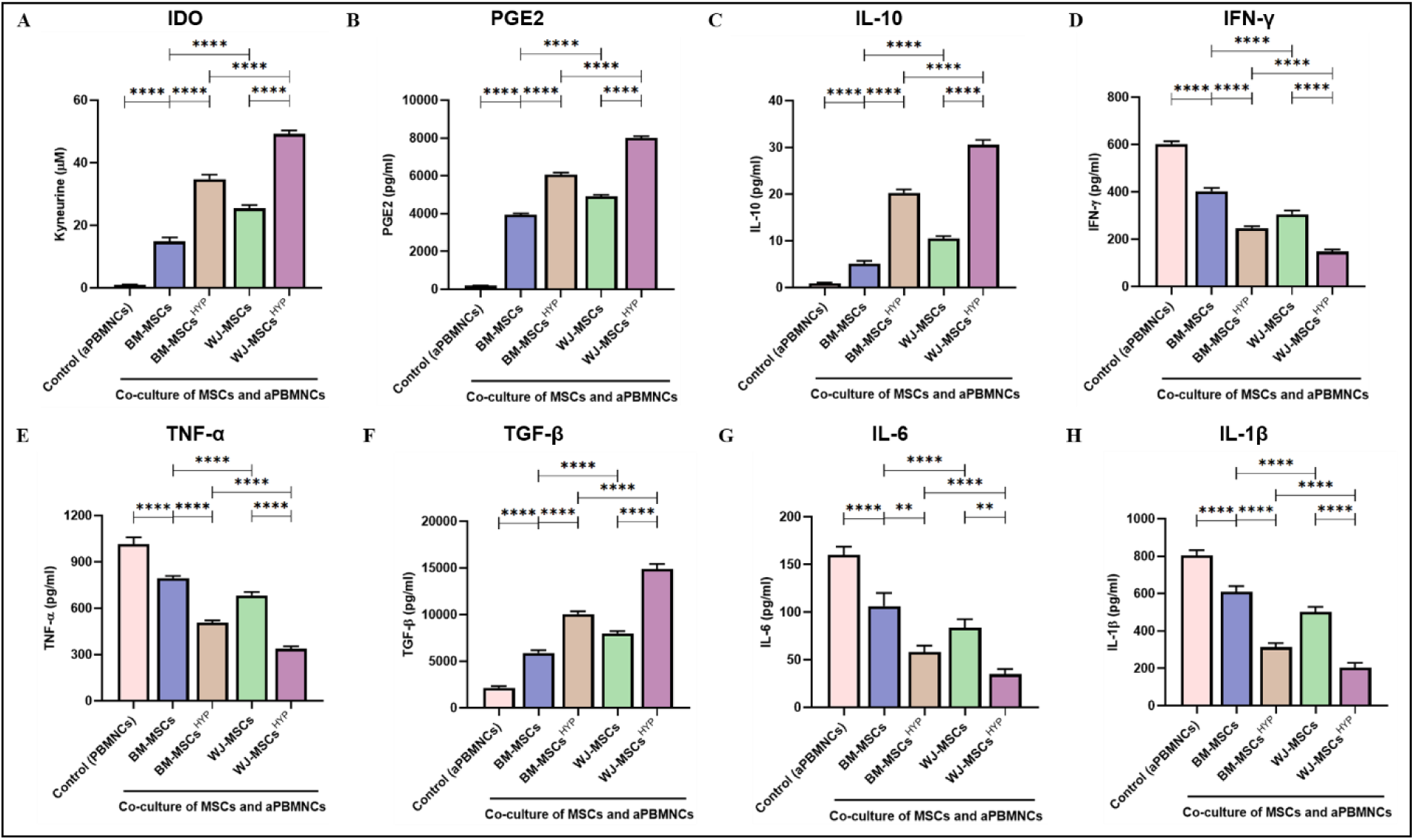
Effect of naïve and hypoxia-preconditioned MSCs (MSCs, MSCs^HYP^) on the secretion of immunomodulatory molecules and cytokines. The bar graphs represent the relative concentration of (A) IDO (μM). (B) PGE2 (pg/ml). (C) IL-10 (pg/ml). (D) IFN-γ (pg/ml). (E) TNF-α (pg/ml). (F) TGF-β (pg/ml). (G) IL-6 (pg/ml). (H) IL-1β (pg/ml) in the direct co-culture of MSCs and aPBMNCs derived from aGVHD patients (n=25). Data shown represent the Mean±S.D of 25 independent experiments performed with PBMNCs derived from 25 different donors (biological replicates), with each experiment conducted in triplicate (technical replicates). Statistical analysis: Tukey’s multiple comparisons test; *≤0.05; **≤0.01; ***≤0.001; ****≤0.0001. *Abbreviations: BM: Bone marrow; WJ: Wharton’s Jelly; MSCs: Mesenchymal stem cells; HYP: Hypoxia-preconditioned; APO: Apoptosis; IDO: Indoleamine 2,3 dioxygenase; PGE2: Prostaglandin E2; IL: Interleukin; IFN-γ: Interferon-γ; TNF-α: Tumor necrosis factor-α; TGF-β: Transforming growth factor-β*

### Apoptosis and efferocytosis in MSCs-mediated immunosuppression and their interaction with CD8⁺ T-cell phenotypes

Our findings demonstrated that both MSCs and MSCs^HYP^ reprogram the immune response of T-cell and APCs under *in vitro* conditions. However, the clinical efficacy of MSCs-based cellular therapy is limited, with approximately 50% of patients responding to treatment (24), and the mechanisms underlying this variability remain poorly understood. Recent studies have suggested that apoptosis of MSCs plays a pivotal role in their therapeutic efficacy (2). This raises the hypothesis that apoptosis of MSCs may be a key determinant of their immunosuppressive efficacy. To explore this, we assessed the apoptosis of MSCs and investigated their correlation with their immunosuppressive effects in a direct co-culture system.

Our morphological observations of the co-culture of MSCs with aPBMNCs revealed that both MSCs and MSCs^HYP^ undergo apoptosis under *in vitro* conditions (Figure S2). Our findings revealed that both MSCs and MSCs^HYP^ undergo apoptosis upon interaction with aPBMNCs followed by efferocytosis. Interestingly, WJ-MSCs exhibited higher apoptosis levels compared to BM-MSCs in both naïve (36.251% vs 27.434%; p: 0.427) and hypoxia-preconditioned conditions (52.60% vs 43.073%; p: 0.357) (Figure 4A), although these differences were not statistically significant. Conversely, BM-MSCs demonstrated higher rates of efferocytosis compared to WJ-MSCs in their naïve (10.001% vs. 5.339; p: 0.595) and hypoxia-preconditioned states (15.813% vs. 10.193; p: 0.4390) (Figure 4B), but these differences were also not statistically significant. Interestingly, efferocytosis exhibited an inverse correlation with apoptosis induction and immune reprogramming potential, where WJ-MSCs consistently outperformed BM-MSCs in these functional parameters.

**Figure 4:**
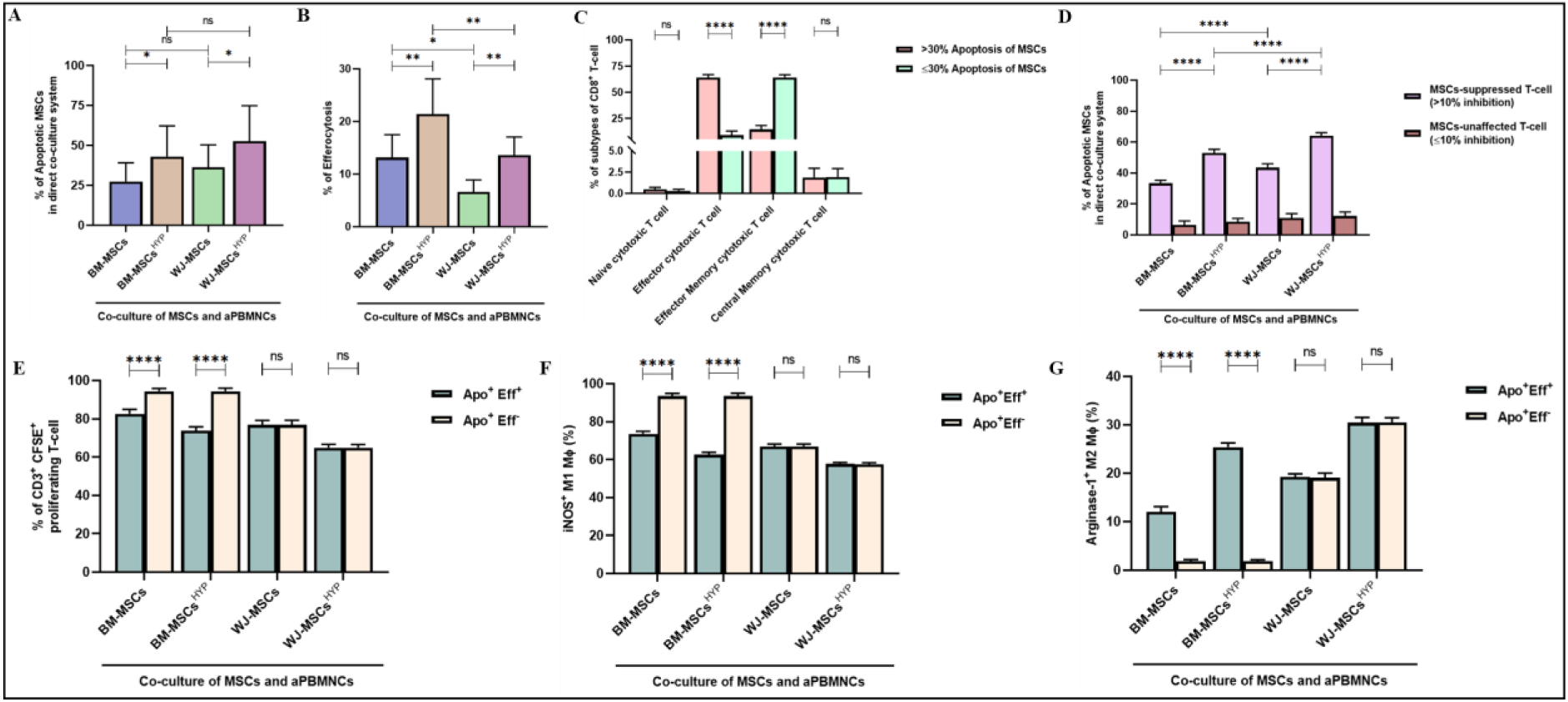
Effect of aGVHD patients derived aPBMNCs on naïve and hypoxia-preconditioned MSCs (MSCs, MSCs^HYP^). The bar graph represents (A) the percentage of apoptotic MSCs (n=25). (B) the percentage of efferocytosis (n=25). (C) the percentage of subtypes of CD8^+^ T-cell (n=25). (D) the percentage of apoptotic MSCs and T-cell suppression (n=25) in the direct co-culture of MSCs and aPBMNCs. Further, the bar graph represents immunomodulation - (E) the percentage of CD3^+^ CFSE^+^ proliferating T-cell (n=25). (F) the percentage of iNOS^+^ M1 macrophage (n=25). (G) the percentage of Arginase-1^+^ M2 macrophage (n=25) under two conditions - one when apoptosis of MSCs followed by their efferocytosis and second one is when MSCs undergo apoptosis only in the direct co-culture of MSCs and aPBMNCs. Data shown represent the Mean±S.D of 25 independent experiments performed with T-cell derived from 25 different donors (biological replicates), with each experiment conducted in triplicate (technical replicates). Statistical analysis: Tukey’s multiple comparisons test; *≤0.05; ****≤0.0001. *Abbreviations: BM: Bone marrow; WJ: Wharton’s Jelly; MSCs: Mesenchymal Stem Cells; HYP: Hypoxia-preconditioned; APO: Apoptosis; Eff: Efferocytosis; aPBMNCs: Activated Peripheral Blood Mononuclear Cells*

These findings prompted us to investigate specific immune cell subsets of aGVHD patients that contributed to the apoptosis of MSCs. To address this, we analyzed the CD4^+^ T-cell and CD8⁺ T-cell subtypes and correlated them with MSCs apoptosis in a direct co-culture system. Our analysis revealed that the effector and effector memory phenotypes of CD8⁺ T-cell were key determinants of MSCs’ fate. In cases where apoptosis exceeded 30%, CD8⁺ T-cell from aGVHD patients predominantly exhibited an effector phenotype (64.10%). In contrast, when MSCs apoptosis was ≤30%, the CD8⁺ T-cell were largely characterized by an effector memory phenotype (64.38%) (Figure 4C).

However, under *in vitro* conditions, no significant variation was observed among CD4^+^ T-cell subtypes with MSCs apoptosis, whether ≤ 30% or ≥ 30%. In both scenarios, effector (50.63%) and effector memory (48.29%) CD4^+^ T-cell phenotypes were the predominant subsets (Figure S3). These findings suggest that CD8^+^ T-cell are the principal mediators of MSCs apoptosis, with CD4^+^ T-cell contributing to a lesser extent. We next evaluated the correlation between MSCs apoptosis and their immunosuppressive potential. Notably, variability was observed in the inhibition of CD3⁺ T-cell proliferation with MSCs apoptosis. Using a median cut-off of 10% inhibition of T-cell proliferation, the study groups were stratified into two categories: ≤10% and >10% inhibition. Interestingly, the group with >10% inhibition demonstrated enhanced MSCs apoptosis, whereas the ≤10% inhibition group exhibited reduced MSC apoptosis across different treatment conditions. Furthermore, WJ-MSCs showed higher levels of apoptosis compared to BM-MSCs (43.482% vs 33.416%; p: ≤0.0001), with hypoxia preconditioning further increasing apoptosis in both MSC types. Despite this, WJ-MSCs^HYP^ exhibited significantly greater apoptosis than BM-MSCs^HYP^ under in vitro co-culture conditions (64.119% vs. 52.932%; *p* ≤ 0.0001) (Figure 4D).

These findings prompt an investigation into whether the immunomodulatory effects observed in aGVHD, including the inhibition of T-cell proliferation, macrophage polarization, and suppression of mature dendritic cells, are driven solely by MSCs apoptosis or require the combined processes of apoptosis and efferocytosis. Specifically, both apoptosis and efferocytosis were necessary to mediate the suppression of T-cell proliferation and the polarization of macrophages towards the M2 phenotype for BM-MSCs and BM-MSCs^HYP^. In contrast, apoptosis alone was the key determinant in suppressing T-cell proliferation, reducing M1 macrophages, and promoting M2 polarization for WJ-MSCs and WJ-MSCs^HYP^ (Figure 4E-G).

### WJ-MSCs^HYP^ elevated mitochondrial transfer and modulated Mitoenergetics and oxidative stress

The intercellular transfer of mitochondria from MSCs has been shown to influence the functional capabilities of immune cells, contributing to the maintenance of immune homeostasis (10). To investigate this phenomenon, we evaluated mitochondrial transfer from MSCs to aGVHD patients-derived activated immune cells and assessed its impact on the immune metabolism under *in vitro* conditions (Figure 5A). Initially, we evaluated the mtDNA content of the MSCs without co-culture, wherein, WJ-MSCs exhibited significantly higher mtDNA copy numbers compared to BM-MSCs (1.1684 vs. 0.722; p ≤0.0001). Hypoxia preconditioning reduced the mtDNA content in both cell types; however, WJ-MSCs^HYP^ retained higher mtDNA levels than BM-MSCs^HYP^ (0.912 vs. 0.516; p ≤0.0001) (Figure 5B).

**Figure 5:**
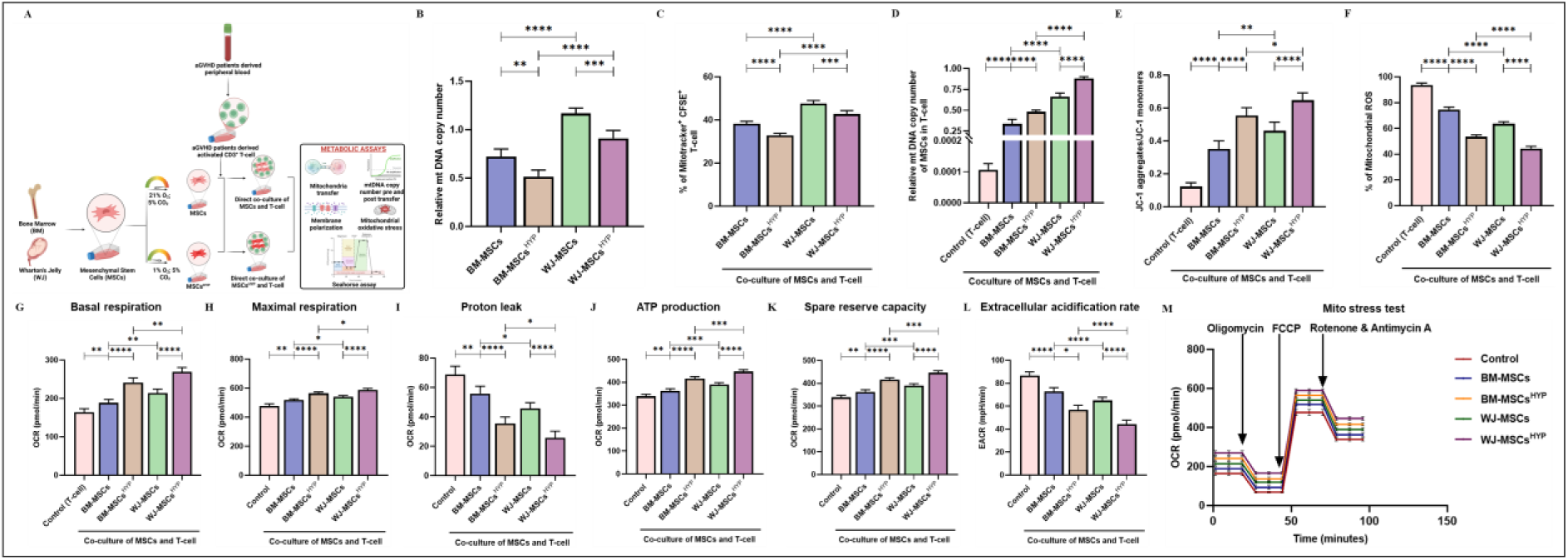
Transfer of mitochondria from MSCs, MSCs^HYP^ modulated T cell metabolism. (A) Diagrammatic representation of the direct co-culture model for the assessment of modulation in T-cell metabolism. The bar graph represents (B) mitochondrial (mt) DNA copy number of MSCs. (C) the percentage of Mitotracker^+^ CFSE^+^ T-cell. (D) mt DNA copy number of MSCs in T-cell post-24-hour co-culture. (E) the ratio of JC-1 aggregates/JC-1 monomers. (F) the percentage of mitochondrial ROS. (G) Basal mitochondrial respiration. (H) Maximal respiration. (I) Proton leak. (J) ATP production. (K) Spare reserve capacity. (L) Extracellular acidification rate. (M) Real-time changes in the oxygen consumption rate of T-cell with or without direct co-culture of MSCs, MSCs^HYP.^ Data shown represent the Mean±S.D of 25 independent experiments performed with T-cell derived from 25 different donors (biological replicates), with each experiment conducted in triplicate (technical replicates). Statistical analysis: Tukey’s multiple comparisons test; *≤0.05; ****≤0.0001. *Abbreviations: BM: Bone marrow; WJ: Wharton’s Jelly; MSCs: Mesenchymal Stem Cells; HYP: Hypoxia-preconditioned; mt: Mitochondria; JC-1:* 5,5′,6,6′-tetrachloro-1,1′,3,3′-tetraethylbenzimidazolocarbocyanine iodide*; OCR: Oxygen consumption rate; EACR: extracellular acidification rate*

Next, we assessed the transfer of mitochondria from MSCs to T-cell and our findings revealed that WJ-MSCs exhibited significantly greater mitochondrial transfer compared to BM-MSCs (47.63% vs. 38.26%; *p* ≤0.0001). Nevertheless, WJ-MSCs^HYP^ maintained a superior transfer efficiency compared to BM-MSCs^HYP^ (42.75% vs. 32.86%; *p* ≤0.0001) (Figure 5C).

Moreover, we quantified the mtDNA content transferred from MSCs to T-cell after co-culture and observed that T-cell co-cultured with WJ-MSCs exhibited significantly higher mtDNA levels compared to those co-cultured with BM-MSCs (0.663 vs. 0.336; p ≤0.0001). Hypoxia preconditioning further enhanced mtDNA transfer, as indicated by the substantial increase in mtDNA content in T-cell co-cultured with MSCs^HYP^. Among these, T-cell co-cultured with WJ-MSCs^HYP^ displayed significantly higher mtDNA levels compared to those co-cultured with BM-MSCs^HYP^ (0.882 vs. 0.481; p ≤0.0001) (Figure 5D).

These findings suggested that mitochondrial transfer may contribute to the modulation of T-cell function and immune metabolism. To address this, we evaluated the mitochondrial health of T-cell from aGVHD patients in the presence or absence of MSCs and MSCs^HYP^ dual staining of mitochondria using JC-1 cationic dye. Cells from aGVHD patients exhibited depolarized mitochondria in their natural microenvironment, as indicated by a reduced JC-1 aggregates/monomers ratio. Both MSCs and MSCs^HYP^ restored mitochondrial polarization, with WJ-MSCs showing a significantly higher JC-1 ratio compared to BM-MSCs (0.462 vs 0.352; p: ≤0.001), and a similar trend was observed in MSCs^HYP^ (0.648 vs. 0.554; *p* ≤0.05) (Figure 5E).

To further assess mitochondrial function, mitochondrial ROS levels were measured using MitoSox dye. Consistent with the mitochondrial polarization data, WJ-MSCs demonstrated greater efficacy in reducing mitochondrial ROS compared to BM-MSCs (63.59% vs. 74.62%; p: ≤0.0001). Hypoxia preconditioning enhanced this effect, with WJ-MSCs^HYP^ exhibiting superior ROS alleviation compared to BM-MSCs^HYP^ (44.46% vs. 53.53%; *p* ≤0.0001) (Figure 5F).

The impact of both MSCs and MSCs^HYP^ on T-cell mitochondrial health was evaluated using the Seahorse Extracellular Flux Mitochondrial Stress Assay. Our findings indicated that both MSCs and MSCs^HYP^, irrespective of their tissue origin, improved the basal respiration rate, maximal respiration rate, ATP production, and spare respiratory capacity, OCR with a concurrent decrease in proton leak and EACR, highlighting the metabolic shift in T-cell. (Figure 5G-M). This shift reflects enhanced mitochondrial health and functional capacity, favoring oxidative phosphorylation over glycolysis, which may contribute to the immunomodulatory effects of MSCs in aGVHD.

### WJ-MSCs enriched in proteins that mediate immune regulation through metabolic reprogramming

In our *in vitro* studies, we observed that WJ-MSCs and WJ-MSCs^HYP^ exhibited superior capabilities in immunomodulation and facilitating the polarization of immune cell mitochondria towards a healthier state compared to BM-MSCs and BM-MSCs^HYP^ respectively. To further elucidate the underlying mechanisms contributing to this enhanced performance, we conducted label-free proteomics analysis on the culture-conditioned media of WJ-MSCs (n=3) alone, as well as in co-culture with aPBMNCs (n=3), alongside separate analyses for BM-MSCs (n=1) and their co-culture with aPBMNCs (n=1).

The comparative analysis of WJ-MSCs in isolation against their co-culture condition revealed a total of 336 dysregulated proteins. Among these, 152 proteins displayed statistically significant alterations with a Log_2_ fold change threshold of ≥ 1.5 and ≤ −1.5. Moreover, we observed that 87 dysregulated proteins were downregulated, while 65 were upregulated in the WJ-MSCs alone and vice versa in the co-culture scenario (Figure 6A-D).

**Figure 6:**
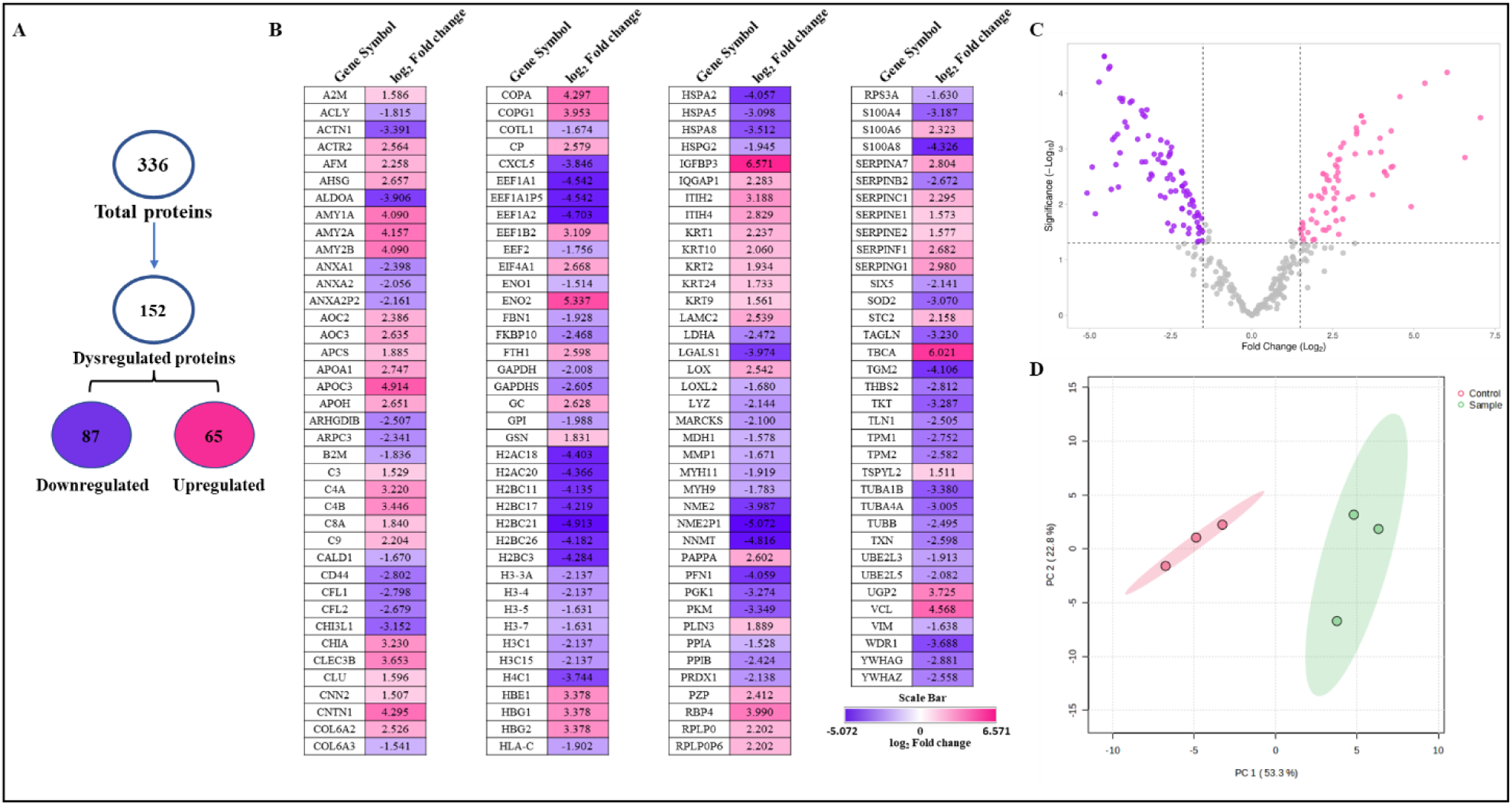
Label-free proteomics analysis of the WJ-MSCs and its direct co-culture with aPBMNCs using LC-MS/MS. (A) A flow chart depicts the total number of identified and dysregulated proteins. (B) A heat map shows the expression levels of dysregulated proteins. (C) A volcano plot highlights differentially expressed proteins. (D) Principal component analysis (PCA) demonstrates the good reproducibility of each biological replicate. Independent experiments were conducted with three different donors (biological replicates).

Gene ontology (GO) analysis of upregulated proteins in WJ-MSCs alone revealed that a significant proportion of the proteins, accounting for 43.75%, were localized within high-density lipoproteins, indicating a noteworthy association with lipid metabolism and transport mechanisms. Following this, 12.73% of the identified proteins were found to reside in the platelet alpha granule lumen, reflecting potential roles in hemostasis and immune regulation. Additionally, 6.25% of the proteins were linked to podosomes, which are specialized structures involved in cell adhesion and matrix degradation, suggesting their involvement in cellular interactions and migration (Figure 7A). Biological and molecular function analyses of the upregulated proteins revealed that WJ-MSCs mediate immune regulation through multiple mechanisms, including complement activation and metabolic reprogramming. Specifically, 30.34% of the enriched proteins were associated with complement activation, with contributions from the classical pathway (18.84%), phagocytosis (5.8%), and endocytosis (1.45%) (Figure 7B-C).

**Figure 7:**
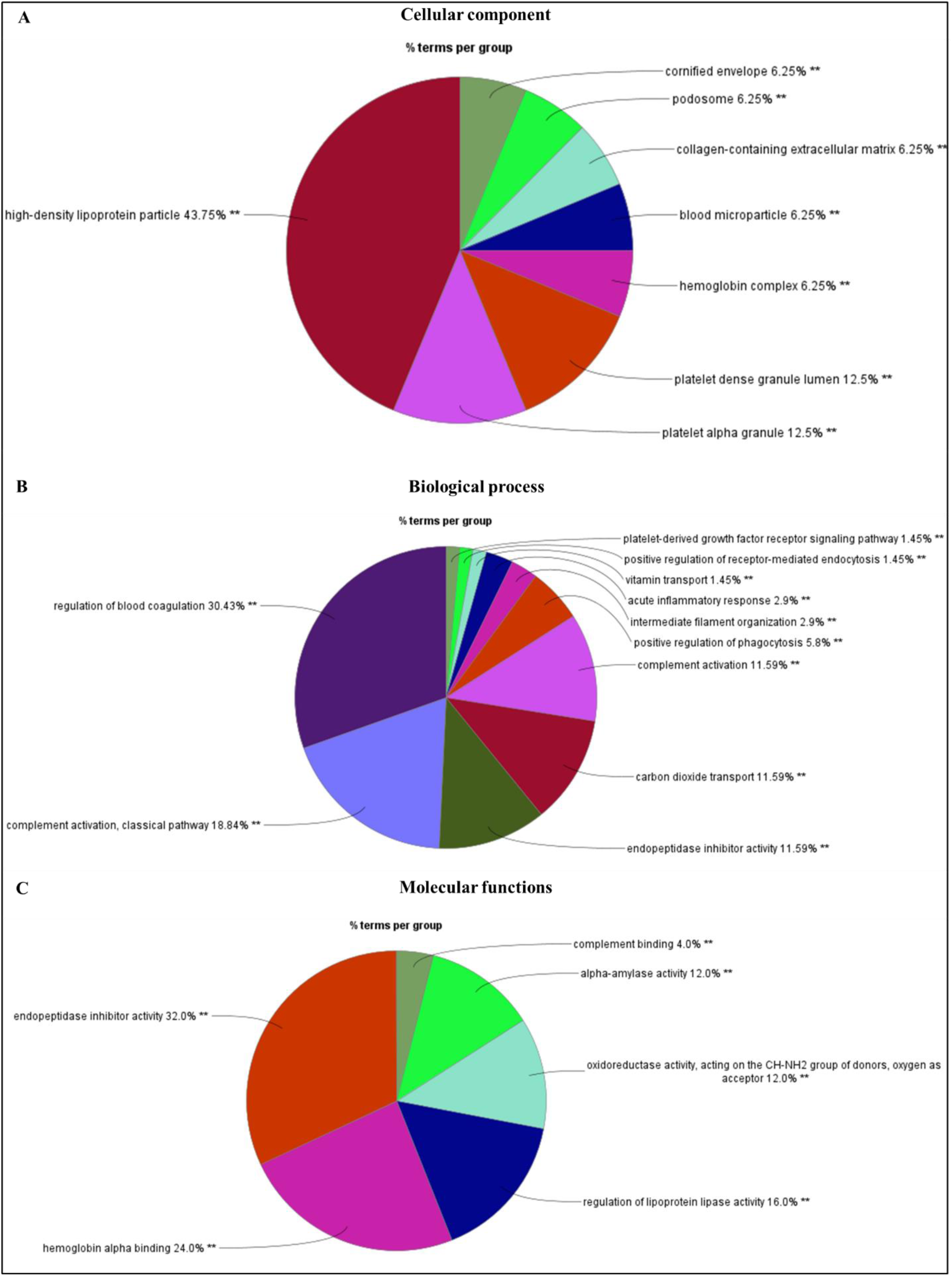
Gene Ontology (GO) analysis of upregulated proteins in WJ-MSCs compared to their co-culture with aPBMNCs. Pie charts depict (A) Cellular component. (B) Biological process. (C) Molecular function. Data showed three independent experiments conducted with three different donors (biological replicates). **≤0.01

Further immune system processes analysis of these upregulated proteins highlighted a significant enrichment of proteins involved in the complement system, including the lectin pathway (42.88%), alternative pathway (14.29%), and classical pathway (28.57%) (Figure 8A-B). KEGG pathway analysis underscored the role of metabolic reprogramming in immune regulation, with significant enrichment of proteins associated with carbohydrate metabolism (33.33%) and cholesterol metabolism (16.67%) (Figure 8C). Reactome pathway analysis further supported these findings, showing the involvement of these proteins in plasma lipoprotein assembly (21.05%), carbohydrate metabolism (15.79%), and complement cascade regulation (26.32%) (Figure 8D).

**Figure 8:**
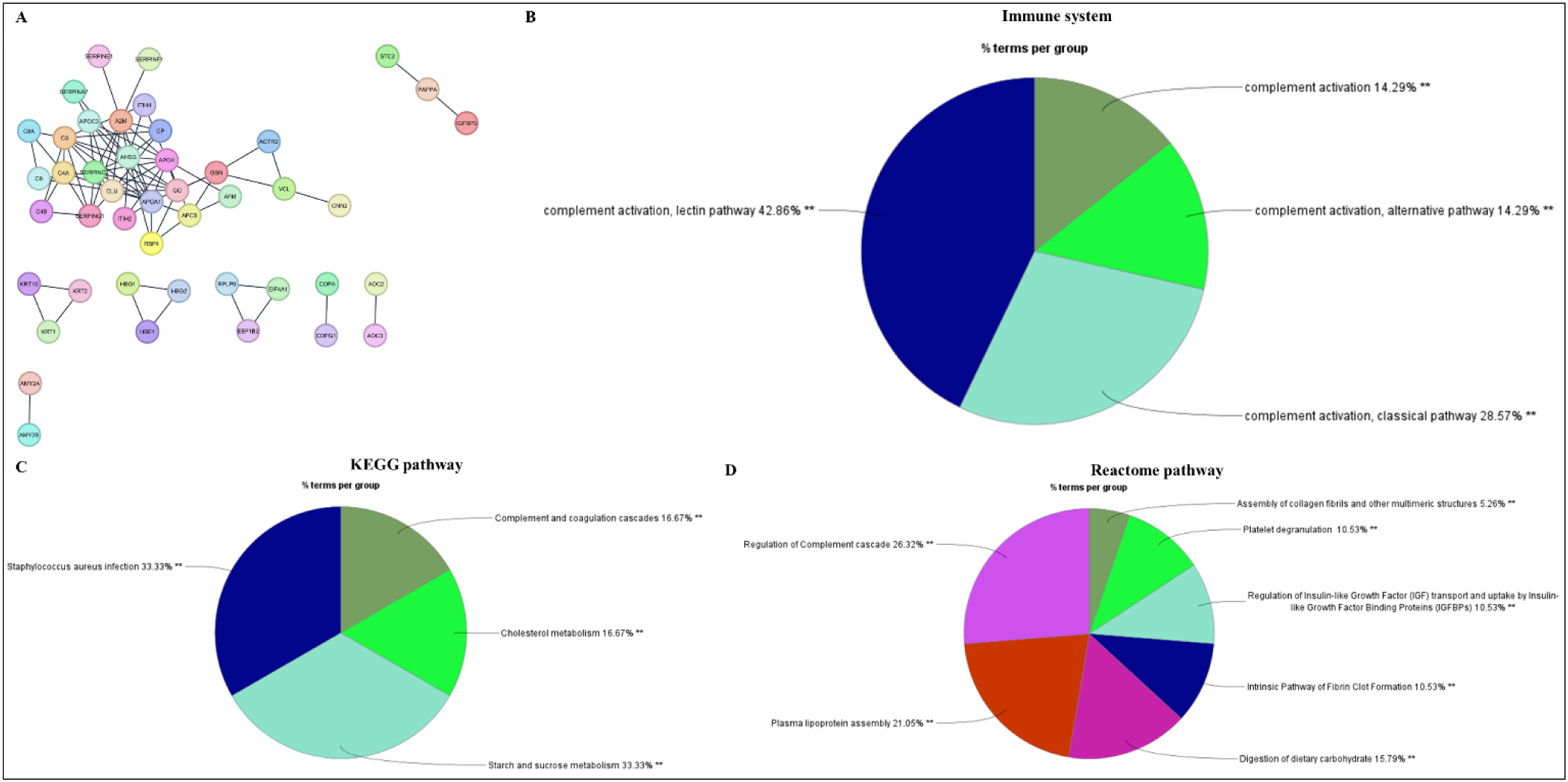
Functional enrichment analysis of upregulated proteins in WJ-MSCs compared to their co-culture with aPBMNCs. (A) STRING network. Pie charts depict (B) Immune system process. (C) KEGG pathway. (D) Reactome pathway. Data showed three independent experiments conducted with three different donors (biological replicates). **≤0.01

Interestingly, the cellular component analysis of the downregulated proteins in the WJ-MSCs alone indicated that 29.41% of the proteins localized to the secretory granule lumen, underscoring a prominent role in the secretion of bioactive molecules. Additionally, 11.76% of the proteins were significantly associated with each cellular structure, including the granule lumen, cortical actin cytoskeleton, focal adhesions, and melanosomes, each contributing to cellular signaling and structural integrity (Figure S4 (I) A). Further analysis of the biological processes demonstrated statistically significant enrichment, with nucleoside diphosphate phosphorylation being the most prevalent, accounting for 35.59% of the downregulated proteins. Other significant biological processes included NADH regeneration (16.69%) and protein refolding (10.17%), indicative of stress response mechanisms involving heat shock proteins (HSPs) and histone proteins (Figure S4 (I) B). Moreover, the analysis showcased various molecular functions, notably significant enrichment in MHC class II protein complex binding (10.53%) and phospholipase A2 inhibitor activity (41.92%), which are critical for immune modulation. Interestingly, 1.69% of the proteins were implicated in the intrinsic apoptotic signaling pathway, while actin filament reorganization (12.04%) and cadherin binding (5.26%) were also statistically significant (Figure S4 (I) C).

The functional enrichment analysis of downregulated proteins in the WJ-MSCs revealed significant insights into immunomodulation and metabolic reprogramming. Notably, 50.0% of the identified proteins were associated with the antimicrobial humoral immune response, specifically mediated by antimicrobial peptides (Figure S4 (II) A-B). In terms of pathway analysis, the KEGG pathways showed that a substantial proportion of the downregulated proteins were involved in systemic lupus erythematosus (SLE) (37.5%), gluconeogenesis (12.5%), and antigen processing and presentation (12.5%) (Figure S4 (II) C). Furthermore, the Reactome pathway analysis indicated that a significant proportion of proteins were involved in the activation of RHO GTPases, which activate PKNs (83.52%), involved in cell growth and differentiation. Additionally, gluconeogenesis (3.3%), alongside gene and protein expression regulated by JAK-STAT signaling following interleukin-12 stimulation (3.3%) was also downregulated in WJ-MSCs alone (Figure S4 (IV) C-D).

The comparative analysis of BM-MSCs in isolation (n=1) against their co-culture condition (n=1) revealed a total of 339 dysregulated proteins displayed statistically significant alterations with a Log_2_ fold change threshold of ≥ 1.5 and ≤ −1.5. Among these, 309 dysregulated proteins were upregulated, while 65 were downregulated in the co-culture of BM-MSCs and aPBMNCs and vice versa in the BM-MSCs alone condition (Figure S5 (I) A-B).

Gene ontology analysis of the upregulated proteins in the co-culture of BM-MSCs and aPBMNCs revealed a shift in the BM-MSC secretory profile towards tissue repair and regeneration. This was characterized by the enrichment of proteins involved in cytoskeleton remodeling (33.74%), integrin-mediated pathways (2.15%), and wound healing (3.76%). Additionally, BM-MSCs showed proteins associated with protection against oxidative stress, including antioxidant activity (4.84%) and thioredoxin peroxidase (3.45%), along with enrichment in proteins involved in proteasome complex formation (5.49%) (Figure S5 (II) A-C). Unlike WJ-MSCs, BM-MSCs exhibited limited involvement in immune regulation, suggesting that BM-MSCs contribute to tissue repair and oxidative stress protection, their immunomodulatory potential and ability to mediate immune regulation are inferior to those of WJ-MSCs. Moreover, the functional analysis revealed that the co-culture of BM-MSCs was enriched in proteins involved in JAK-STAT signaling through IL-12 stimulation (6.25%) and neutrophil degranulation and its apoptotic processes (15.78%). However, a notable enrichment of proteins associated with viral carcinogenesis (40.0%), as well as infections caused by *Salmonella* (10.0%) and *E. coli* (6.67%), was observed. This suggests a potential vulnerability of BM-MSCs to secondary infections, further limiting their suitability for robust immune modulation compared to WJ-MSCs (Figure S5 (III) A-D).

## DISCUSSION

Recent studies have shown that hypoxia preconditioning not only mimics the MSCs’ native niche but also optimizes their therapeutic potential by enhancing their resilience to oxidative stress and inflammation. In this context, our study elucidates how the hypoxic microenvironment of MSCs enhances their immunomodulatory functions and optimizes mitochondrial health in immune cells derived from aGVHD patients.

Our study demonstrated that hypoxia-preconditioned MSCs retained their fibroblast-like, spindle-shaped morphology, surface profile, and trilineage differentiation potential, regardless of tissue origin. These findings are consistent with our previous work and that of others (4,25–28). However, a few studies reported that hypoxia preconditioning decreased their potential to differentiate into adipogenic and osteogenic (29) with an increase in their potential to be chondrogenic (30).

Moreover, the metaphase analysis of both BM- and WJ-MSCs demonstrated a normal diploid karyotype, with no detectable chromosomal aberrations upon G-banding, indicating that the MSCs exhibited chromosomal stability. These results are consistent with previous studies, which have shown that MSCs isolated from both adult and fetal tissue sources maintain chromosomal integrity, making them suitable for subsequent applications (31–34).

Recent studies investigating the impact of hypoxia preconditioning on MSCs have provided valuable insights into their immunomodulatory properties, particularly in modulating T-cell and macrophage responses. Our findings demonstrated that 1% O₂ preconditioning significantly enhanced the immunoregulatory effects of MSCs. This was evident by a substantial reduction in CD3^+^ T-cell proliferation, increased induction of Tregs, and polarization of CD4^+^ T-cell towards Th2 and Th9 phenotypes, alongside a suppression of Th17 responses. Notably, WJ-MSCs^HYP^ outperformed BM-MSCs^HYP^ under *in vitro* conditions, underscoring their potential to maintain immune tolerance and promote an environment favorable for humoral immunity and tissue repair. These results align with our own and other studies showing that preconditioning of umbilical cord/umbilical cord-blood-derived MSCs with either 1% O₂ (4,35,36), 2% O₂ (37), or 3% O_2_ (38) alleviate inflammation by decreasing T-cell proliferation.

To the best of our knowledge, none of the prior studies have investigated the effect of naive and hypoxia-preconditioned MSCs on the γδ and αβ proportion of CD4^+^ T-cell. In our study, we observed 1% O_2_ selectively reduced the proportion of αβ CD4^+^ T-cell in aGVHD, suggesting differential regulatory effects on T-cell subsets and could be attributed to enhanced differentiation towards Tregs. Previous studies revealed that the maintenance or slight increase in γδ CD4^+^ T-cell can be indicative of a shift towards a more regulatory immune environment that can modulate inflammation, and promote immune homeostasis and tissue repair (39). For instance, MSCs have been shown to enhance the conversion of pro-inflammatory Th17 into Tregs, which may further explain the reduction in αβ CD4^+^ T-cell populations (40,41).

In line with the findings of T-cell, WJ-MSCs^HYP^ significantly enhanced the polarization of M1 macrophage to anti-inflammatory M2 phenotype, consistent with our previous findings (4).

Available literature has shown that the immunoregulatory effects of MSCs are primarily mediated through the paracrine mechanism (42–44). Similarly, we observed that MSCs^HYP^ significantly increased levels of IDO, PGE2, IL-10, and TGF-β with a concurrent decrease in pro-inflammatory cytokines such as IL-6, IFN-γ, TNF-α, and IL-1β. These findings corroborate their immunomodulatory effects on the aGVHD patients-derived immune cells in a cellular model. Notably, the observed increase in IDO and PGE2 levels correlates with the inhibition of T-cell proliferation and the induction of Tregs, caused by a rise in TGF-β. This modulation is further supported by the concurrent rise in IL-10 and TGF-β, which are known to facilitate M2 macrophage polarization. The decrease in IFN-γ and TNF-α levels contributes to a reduction in M1 macrophage activation, thereby shifting the immune response toward a more anti-inflammatory state. Additionally, the reduction in IL-6 and IL-1β corresponds with a decrease in Th17 and Th1 phenotypes, respectively.

Recent studies have provided compelling evidence that MSCs, particularly when administered *in vivo*, tend to undergo apoptosis, a process that has been found to facilitate their immunomodulatory effects. In our study, we observed that WJ-MSCs^HYP^ exhibits a higher propensity for apoptosis compared to BM-MSCs^HYP^ when comes in contact with immune cells, consistent with our previous findings (4). The differential susceptibility of these two MSCs sources to apoptosis may allow for the development of more effective treatment regimens tailored to specific inflammatory diseases. Previous studies have predominantly focused on BM-MSCs, which serve as the conventional reference point for evaluating the therapeutic capabilities of MSCs in various immune-related disorders (2).

Studies revealed that the apoptosis of BM-MSCs is a pivotal mechanism influencing their immunosuppressive effects; upon cell death, these MSCs can be efficiently phagocytosed by macrophages, leading to an altered immune response (3). Such immunological programming via efferocytosis suggests that apoptotic BM-MSCs can modulate inflammation through the secretion of bioactive molecules that contribute to an anti-inflammatory environment. Interestingly, our study has yielded notable findings regarding the efferocytosis of MSCs, revealing a distinct contrast between BM-MSCs and WJ-MSCs in both normoxic and hypoxic conditions. Specifically, we found that BM-MSCs exhibit a greater capacity for efferocytosis compared to their WJ-MSC counterparts, similar to our previous findings (4). This observation presents an intriguing paradox, especially in light of our previous findings highlighting that WJ-MSCs undergo apoptosis at a higher rate than BM-MSCs. However, none of the studies explored efferocytosis of apoptotic WJ-MSCs.

In our study, effector phenotypes of CTLs predominantly drive MSC apoptosis, significantly influencing the immunosuppressive capacity of MSCs. This aligns with previous findings suggesting that the activation state of CTLs is crucial for the modulation of MSCs function. Specifically, there is evidence demonstrating that a heightened presence of effector CTLs correlates with increased apoptosis of MSCs, thereby enhancing the immunosuppressive effects on CD3^+^ T-cell, which are vital for adaptive immune responses (45). This phenomenon may be attributed to the fact that activated CTLs can induce MSCs death through perforin and granzyme-mediated pathways, which are known to lead to rapid cellular demise and subsequent immunological implications (2). The differential impact of CTL subsets (effector, effector memory) on MSCs viability indicates a nuanced interplay between T-cell activation state and MSCs-mediated immunosuppression. It has been suggested that effector memory CTLs may engage in a more regulated interaction with MSCs, potentially limiting pro-inflammatory signals that could lead to MSCs apoptosis. In contrast, effector CTLs evoke a more aggressive immune response, promoting MSCs death and enhancing the immunosuppressive environment created by the dying MSCs.

Our investigation into the interactions between BM-MSCs and WJ-MSCs with immune cells reveals important distinctions in their mechanisms of immunosuppression, particularly in the context of apoptosis and efferocytosis. We observed that concomitant with apoptosis, BM-MSCs effectively inhibited T-cell proliferation, which aligned with our previous work and documented the ability of MSCs to create an immunosuppressive environment following their apoptosis (4). This mechanism is reinforced by the subsequent efferocytosis of apoptotic BM-MSCs by macrophages, facilitating their polarization towards anti-inflammatory phenotypes. In contrast to the role of BM-MSCs, our findings indicate that the immunosuppressive effects mediated by WJ-MSCs do not significantly depend on efferocytosis. While apoptosis of WJ-MSCs does lead to some level of inhibition in T-cell proliferation, it appears that the efferocytosis process does not serve as a crucial determinant in augmenting their immunosuppressive potential. This lack of reliance on efferocytosis may indicate that WJ-MSCs employ alternative mechanisms to exert their immunosuppressive effects, such as the direct secretion of soluble factors that modulate immune responses, independent of the engulfment of apoptotic debris. To the best of our knowledge, this is the first study that explored the differential mechanism of immunosuppression mediated by BM-MSCs and WJ-MSCs.

The role of mitochondria in mediating immune suppression through MSCs has garnered increasing attention, in this context, our study highlights significant distinctions in mitochondrial dynamics between tissue-specific human MSCs, with WJ-MSCs demonstrating a notably higher mtDNA content compared to their BM counterparts. This finding aligns with previous research indicating that mtDNA levels can directly correlate with metabolic activity (46) and immunoregulatory potential of stem cells (47).

Interestingly, we found that hypoxia preconditioning led to a decrease in mtDNA content across both MSCs, suggesting a universal response to low oxygen conditions that might impact their metabolic pathways. However, it is noteworthy that despite this reduction in mtDNA levels, hypoxia preconditioning did not impede the mitochondrial transfer capability of either WJ-MSCs or BM-MSCs, with over 95% of mitochondria successfully transferred to T-cell, as evidenced by increased mtDNA content in these recipient cells. This is consistent with findings from previous studies, which affirm that mitochondrial transfer plays a critical role in modulating immune cells during inflammation and tissue repair processes (48–50).

Moreover, our study demonstrated that WJ-MSCs^HYP^ exhibits superior efficiency in polarizing T-cell mitochondria to a polarized state compared to BM-MSCs^HYP^. The ability of WJ-MSCs^HYP^ to mitigate mitochondrial ROS highlights their potential in restoring metabolic homeostasis within T-cell, thereby facilitating improved immune responses. This adaptation appears to alleviate mitochondrial ROS levels in T-cell, bridging the link between mitochondrial health and immune cell functionality (51).

Our findings confirm that this transfer significantly enhances various aspects of T-cell metabolism and functionality. Specifically, we observed an increase in basal respiration, maximal respiration, ATP production, and OCR in T-cell following mitochondrial transfer. These enhancements indicate a robust metabolic shift towards improved bioenergetics, aligning with the literature that has demonstrated the role of MSC-derived mitochondria in bolstering T-cell function and survival in inflammatory contexts (48).

Our data also revealed an increase in spare respiratory capacity, suggesting that T cells receiving mitochondria from MSCs are better equipped to respond to fluctuations in energy demand during activation. The concurrent decrease in EACR and proton leak further underscores the improved metabolic efficiency and mitochondrial health of T cells post-transfer. This is particularly noteworthy since excessive proton leak is often associated with mitochondrial dysfunction and increased production of ROS, which can lead to cellular senescence and impaired immune responses (50).

Notably, the heightened mitochondrial performance in T-cell was especially pronounced in those interacting with WJ-MSCs^HYP^. This observation suggests that WJ-MSCs^HYP^ not only facilitate effective mitochondrial transfer but also possibly undergo metabolic adaptations under hypoxic conditions that enhance their mitochondrial content and functionality. Previous studies have indicated that hypoxic environments can prime MSCs for improved therapeutic efficacy, potentially leading to enhanced mitochondrial integrity and functional capacity when transferred to recipient immune cells (37,51).

Our study illustrates the nuanced roles of mitochondrial transfer, apoptosis, and efferocytosis in immune regulation. Mitochondrial transfer and apoptosis of WJ-MSCs significantly contribute to immunosuppressive effects, independent of reliance on efferocytosis, and offer new insights into their therapeutic utility. In contrast, mitochondrial transfer in conjunction with apoptosis and subsequent efferocytosis underscores a multifaceted mechanism of action in immune modulation by BM-MSCs. This complexity reflects the inherent capabilities of different MSCs sources to alleviate inflammation and create a homeostatic microenvironment in aGVHD.

Our proteomic analyses revealed that WJ-MSCs mediate immune regulation through the involvement of the complement system, lipid metabolism, and carbohydrate metabolism. This aligns with previous studies suggesting that MSCs can modulate T-cell responses through various mechanisms, including the activation of complement cascades and the secretion of immunoregulatory factors such as indoleamine 2,3-dioxygenase (IDO) and hepatocyte growth factor (HGF) (52,53).

In contrast, the comparative analysis of BM-MSCs revealed a predominance of proteins involved in tissue repair and oxidative stress protection rather than direct immune modulation. Although BM-MSCs exhibited a robust response to oxidative stress through antioxidant activity and thioredoxin peroxidase enrichment, their limited immunomodulatory capabilities highlight a functional divergence between BM-MSCs and WJ-MSCs. This is consistent with findings that suggest WJ-MSCs possess enhanced immunosuppressive properties compared to BM-MSCs, likely due to their distinct secretory profiles and gene expression patterns (54,55). Additionally, when BM-MSCs come into contact with immune cells, they are enriched in proteins that promote JAK-STAT signaling and predominantly target neutrophil activity. This makes them less suitable for aGVHD therapy, as aGVHD is characterized by the activation of JAK-STAT signaling and primarily driven by T-cell-mediated immune responses (56). These findings make BM-MSCs less effective for aGVHD, as it leads to an imbalance that favors inflammation over resolution.

Despite the significant findings presented in this study regarding the immunomodulatory roles of hypoxia-preconditioned MSCs, several limitations must be acknowledged. First, the study primarily focuses on in vitro models which, while valuable, may not fully replicate the complexities of the in vivo environment. The interactions between MSCs and immune cells, as well as the subsequent functional outcomes, can greatly be influenced by factors such as the tissue microenvironment, systemic cytokine profiles, and underlying disease states, which are challenging to mimic in laboratory conditions.

Second, while we explored the differential effects of BM-MSCs and WJ-MSCs on various T-cell subsets, our investigation did not encompass a comprehensive analysis of all immune cell types that could be influenced by these MSCs, such as B cells or dendritic cells. Future studies should consider the broader immune landscape to fully understand the systemic impacts of MSCs therapies.

## CONCLUSION

The potential for WJ-MSCs^HYP^ to optimally restore T-cell functionality while simultaneously promoting anti-inflammatory responses underscores their therapeutic promise in treating inflammatory diseases, including aGVHD. Further investigations into the molecular underpinnings of these mechanisms and the translational potential of MSCs-based therapies could pave the way for innovative treatment strategies aimed at restoring immune homeostasis and improving patient outcomes.

## ABBREVIATIONS

MSCs: Mesenchymal Stem Cells
aGVHD: Acute Graft-versus-Host-Disease
BM: Bone marrow
WJ: Wharton’s Jelly
HYP: Hypoxia
HIF: Hypoxia Inducible Factors
ISCT: International Society for Cellular Therapy
DMEM: Dulbecco’s Modified Eagle Medium
RPMI: Rosewell Park’s Memorial Institute
aPBMNCs: Activated Peripheral Blood Mononuclear Cells
CFSE: Carboxyfluorescein diacetate succinimidyl ester
JC1: 5,5′,6,6′-tetrachloro-1,1′,3,3′-tetraethylbenzimi-dazolylcarbocyanine iodide
Th: Helper T-cell
Tc: Cytotoxic T-cell
CTLs: Cytotoxic T lymphocytes
IL: Interleukin
IDO: Indoleamine 2,3-dioxygenase
PGE2: Prostaglandin E2
TNF-α: Tumor Necrosis Factor-alpha
TGF-β: Transforming Growth Factor-beta
IFN-γ: Interferon-gamma
mtDNA: Mitochondria DNA
ROS: Reactive Oxygen Species
OCR: Oxygen Consumption Rate
EACR: Extracellular Acidification Rate

## ACKNOWLEDGEMENT

The authors express their gratitude to the All India Institute of Medical Sciences (AIIMS), New Delhi, India for facilitating the execution of the study. Schematic representative figures illustrating the methodology were created using Biorender.com.

## AUTHOR’S CONTRIBUTIONS

MM performed the experiments, acquired, analyzed, and interpreted the data, and wrote the manuscript. MM and VS were involved in performing experiments and data interpretation and analysis. LM was involved in the interpretation of proteomics results and drafted the proteomics section of the manuscript. SR and RG contributed to data interpretation and analysis. PD was involved in editing of the manuscript. SB, VD, DP, MA, AKG, PSM, RP, MM, TS, and RD provided patient samples and their clinical details. SB and GH were involved in the interpretation of proteomics data. BN, TDS, SK, RAM, and SKS contributed to data interpretation and analysis. HP, SM, and RKS conceptualized the study, designed and supervised the experiments, interpreted the data, and reviewed and edited the manuscript. All authors critically reviewed, and approved the final version of the manuscript.

---

**Figure S1:**
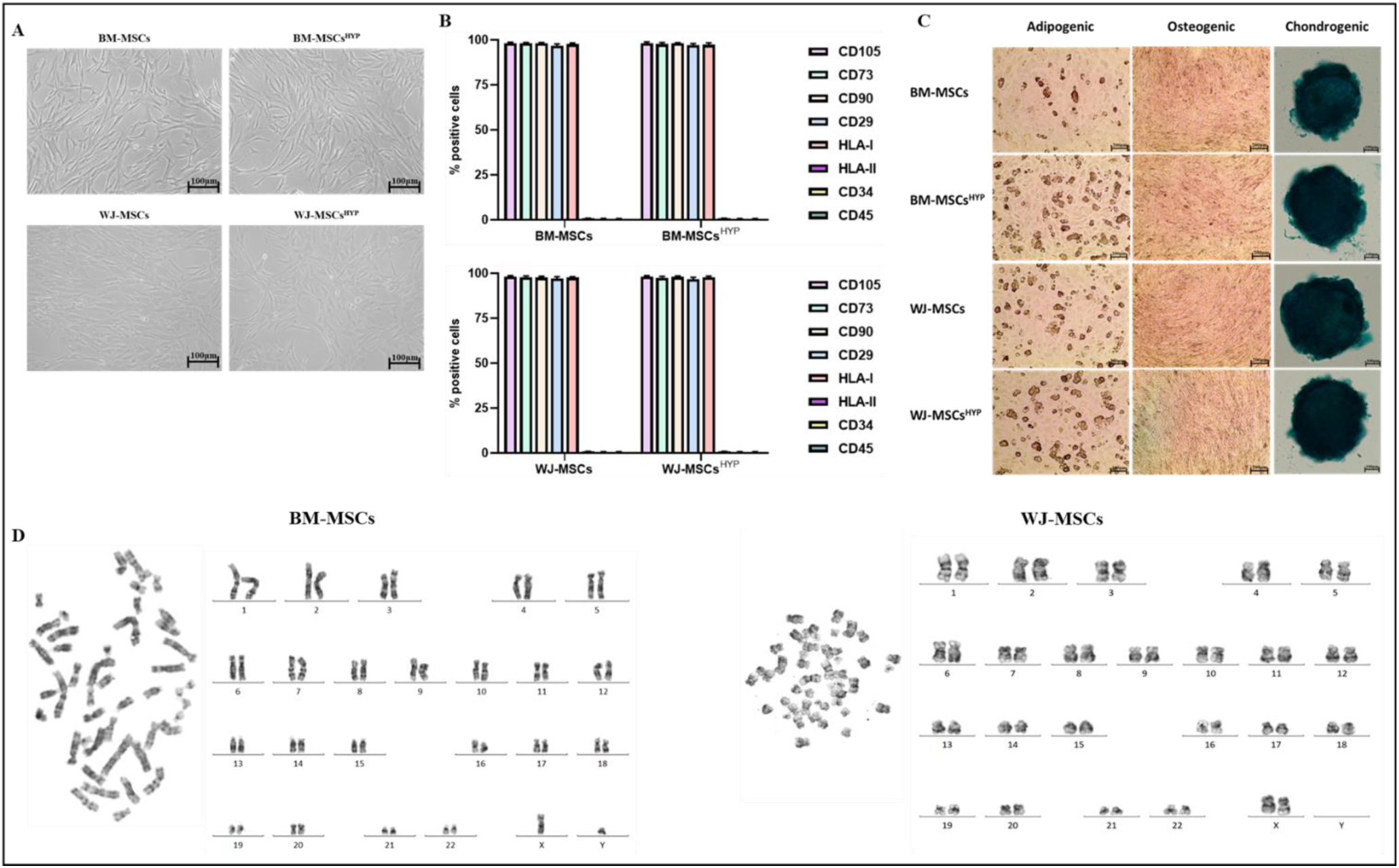
Characterization of tissue-specific human MSCs (BM-MSCs, BM-MSCs^HYP^, WJ-MSCs, and WJ-MSCs^HYP^) **(A)** Morphological images. **(B)** Bar graphs depict surface marker profiling using flow cytometry. **(C)** Trilineage differentiation. Data are shown as Mean±S.D. The data shown are from independent experiments performed with MSCs derived from three different donors (biological replicates) and conducted in triplicates (technical replicates).

**Figure S2:**
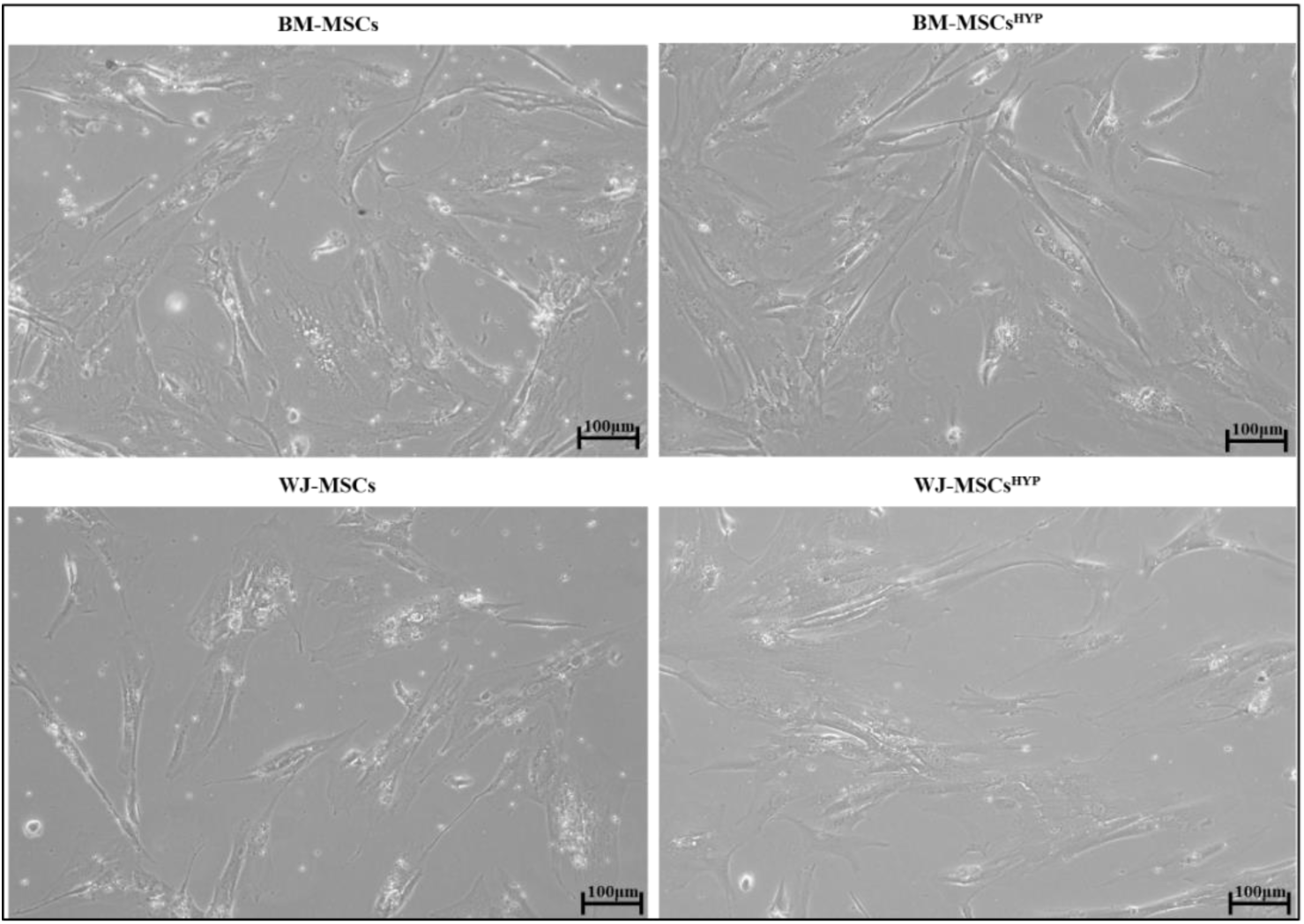
Morphological images of direct co-culture of tissue-specific human MSCs (BM-MSCs, BM-MSCs^HYP^, WJ-MSCs, and WJ-MSCs^HYP^) and aGVHD patients derived T-cell. *Abbreviations: BM: Bone marrow; WJ: Wharton’s Jelly; MSCs: Mesenchymal Stem Cells; HYP: Hypoxia-preconditioned;*

**Figure S3:**
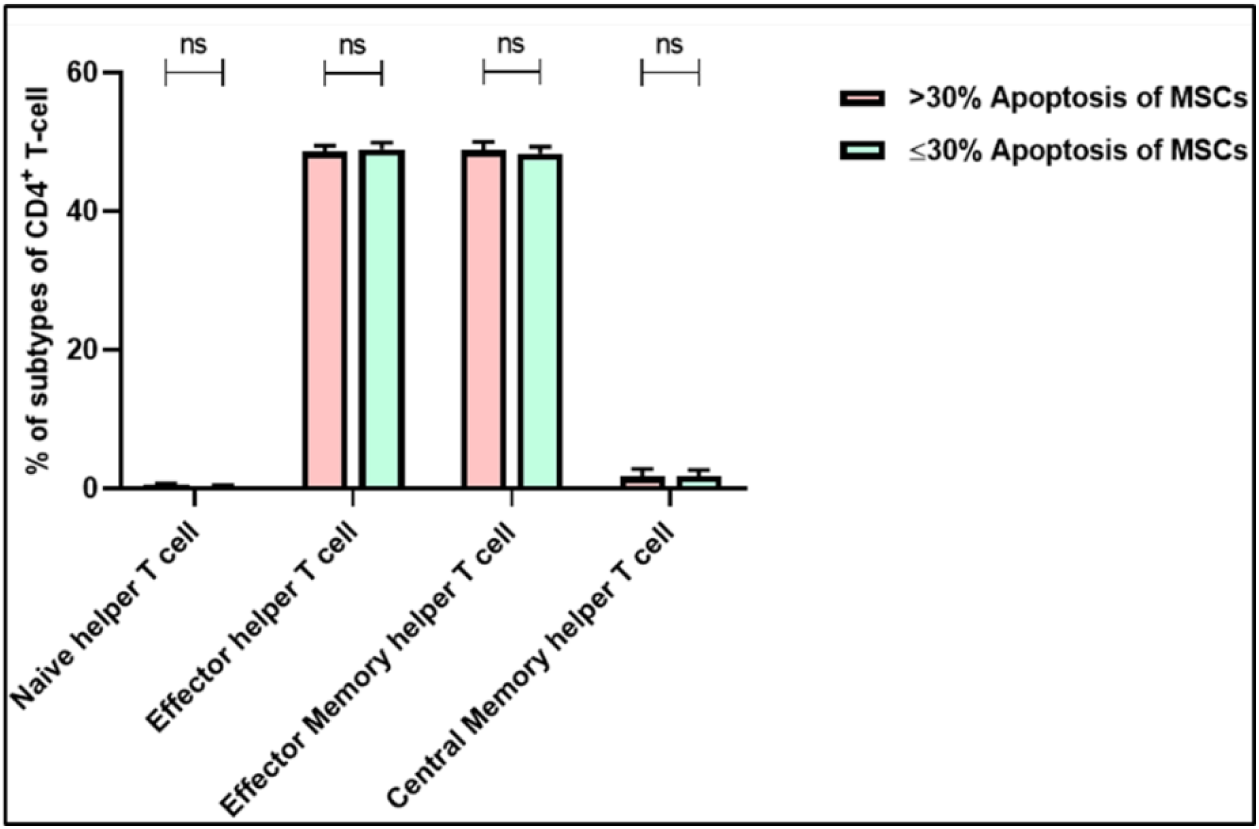
The bar graph represents the percentage of subtypes of CD4^+^ T-cell (n=25). Data shown represent the Mean±S.D of 25 independent experiments performed with T-cell derived from 25 different donors (biological replicates), with each experiment conducted in triplicate (technical replicates). Statistical analysis: Tukey’s multiple comparisons test; *≤0.05; ****≤0.0001. *Abbreviations: BM: Bone marrow; WJ: Wharton’s Jelly; MSCs: Mesenchymal Stem Cells; HYP: Hypoxia-preconditioned; APO: Apoptosis; Eff: Efferocytosis; aPBMNCs: Activated Peripheral Blood Mononuclear Cells*

**Figure S4 (I):**
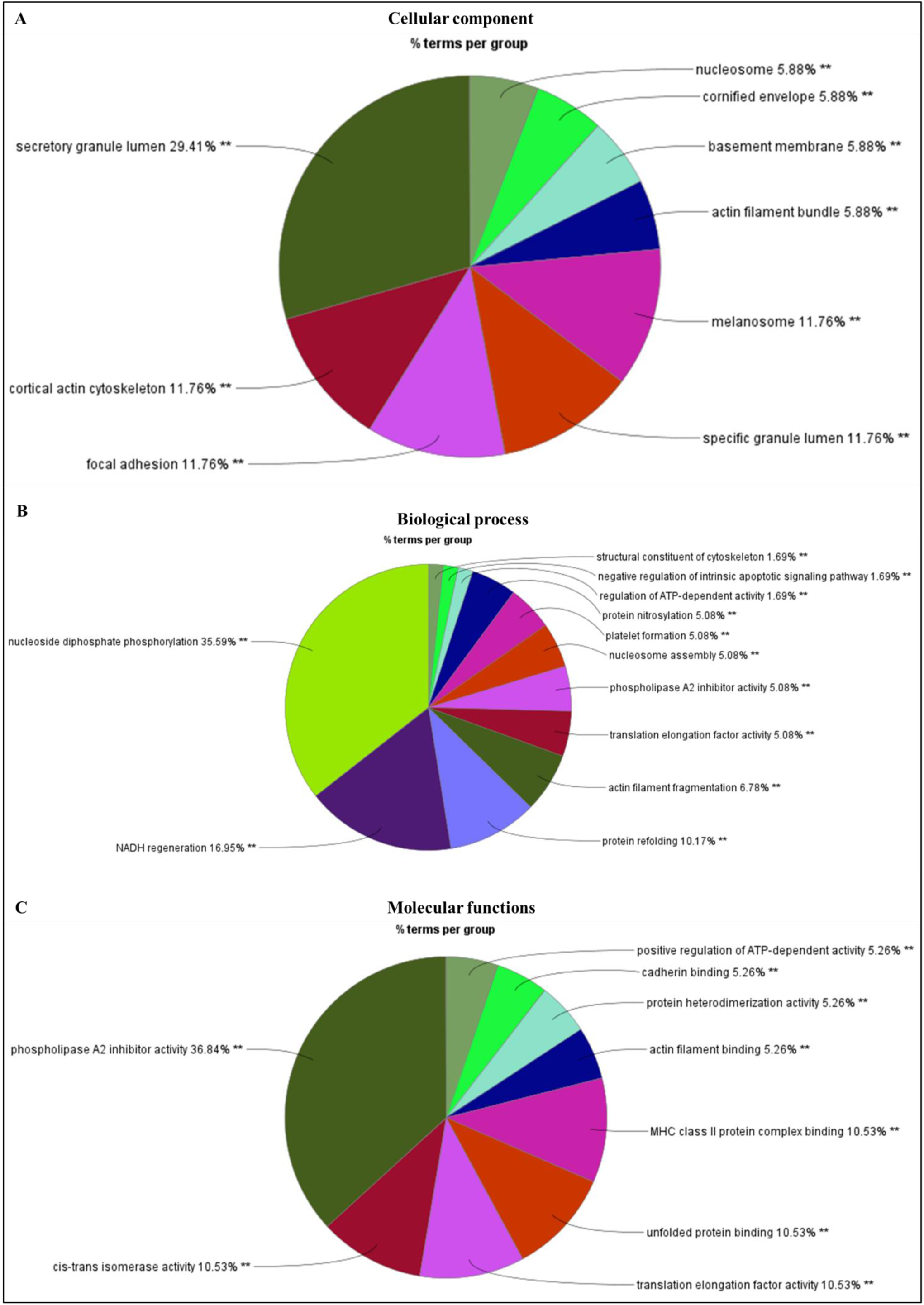
Gene Ontology (GO) analysis of downregulated proteins in WJ-MSCs compared to its co-culture with aPBMNCs. Pie charts depict (A) Cellular component. (B) Biological process. (C) Molecular function. Data showed three independent experiments conducted with three different donors (biological replicates).

**Figure S4 (II):**
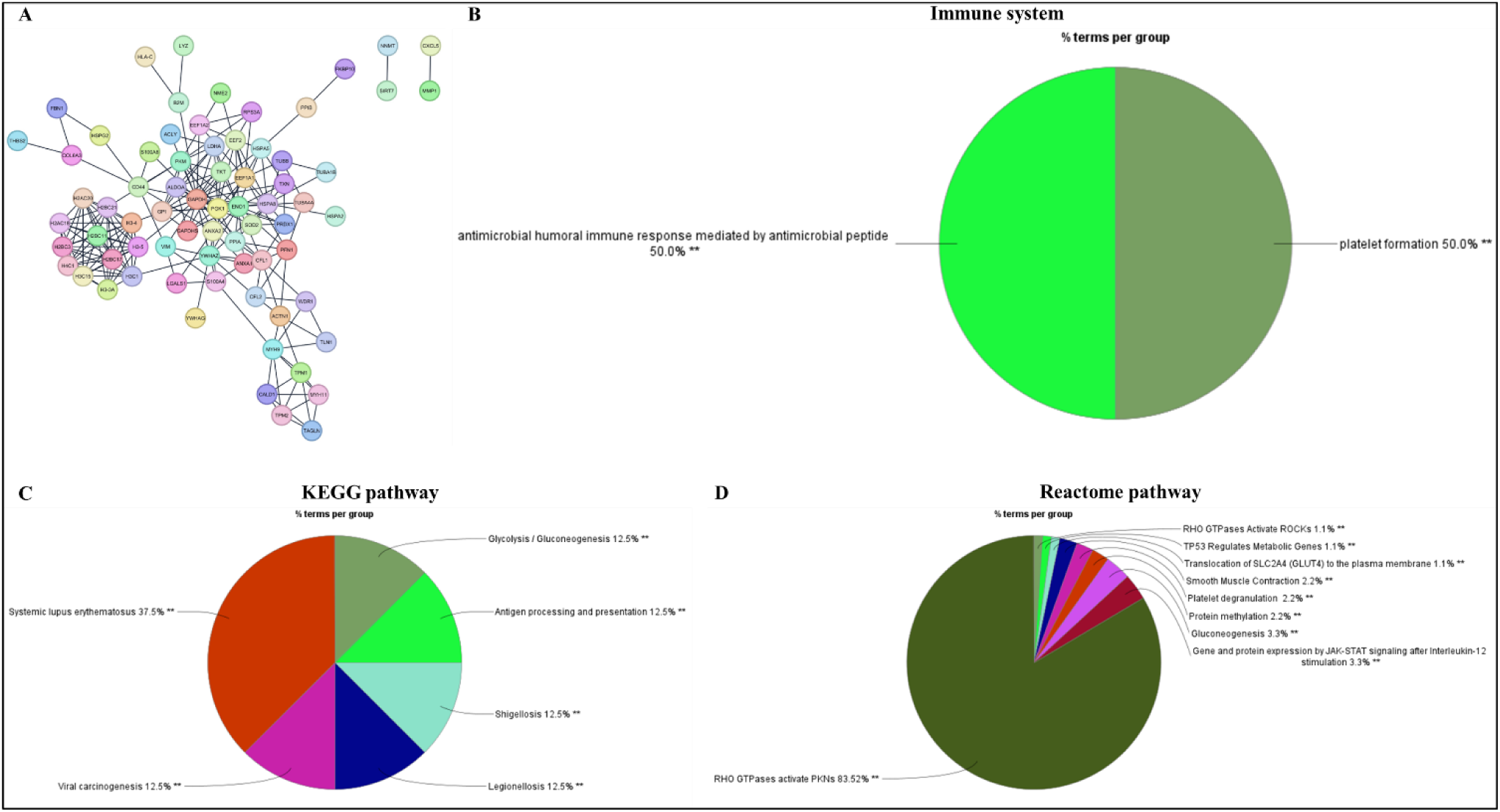
Functional enrichment analysis of downregulated proteins in WJ-MSCs compared to its co-culture with aPBMNCs. (A) STRING network. Pie charts depict (B) Immune system process. (C) KEGG pathway. (D) Reactome pathway. Data showed three independent experiments conducted with three different donors (biological replicates).

**Figure S5:**
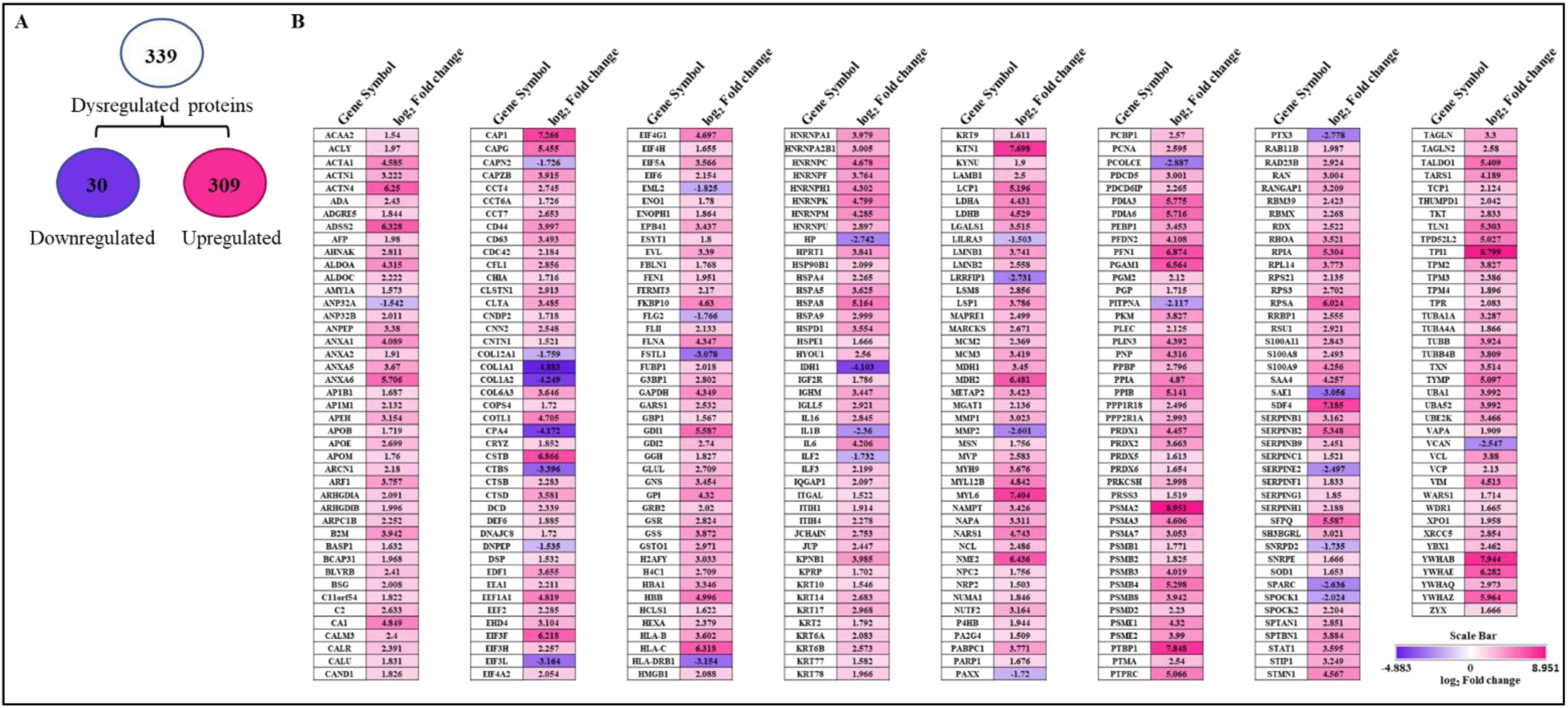
Label-free proteomics analysis of BM-MSCs and its coculture with aPBMNCs using LC-MS/MS. (A) A flow chart depicts the total number of identified and dysregulated proteins. (B) A heat map shows the expression levels of dysregulated proteins. An experiment was conducted with a single donor.

**Figure S5 (II):**
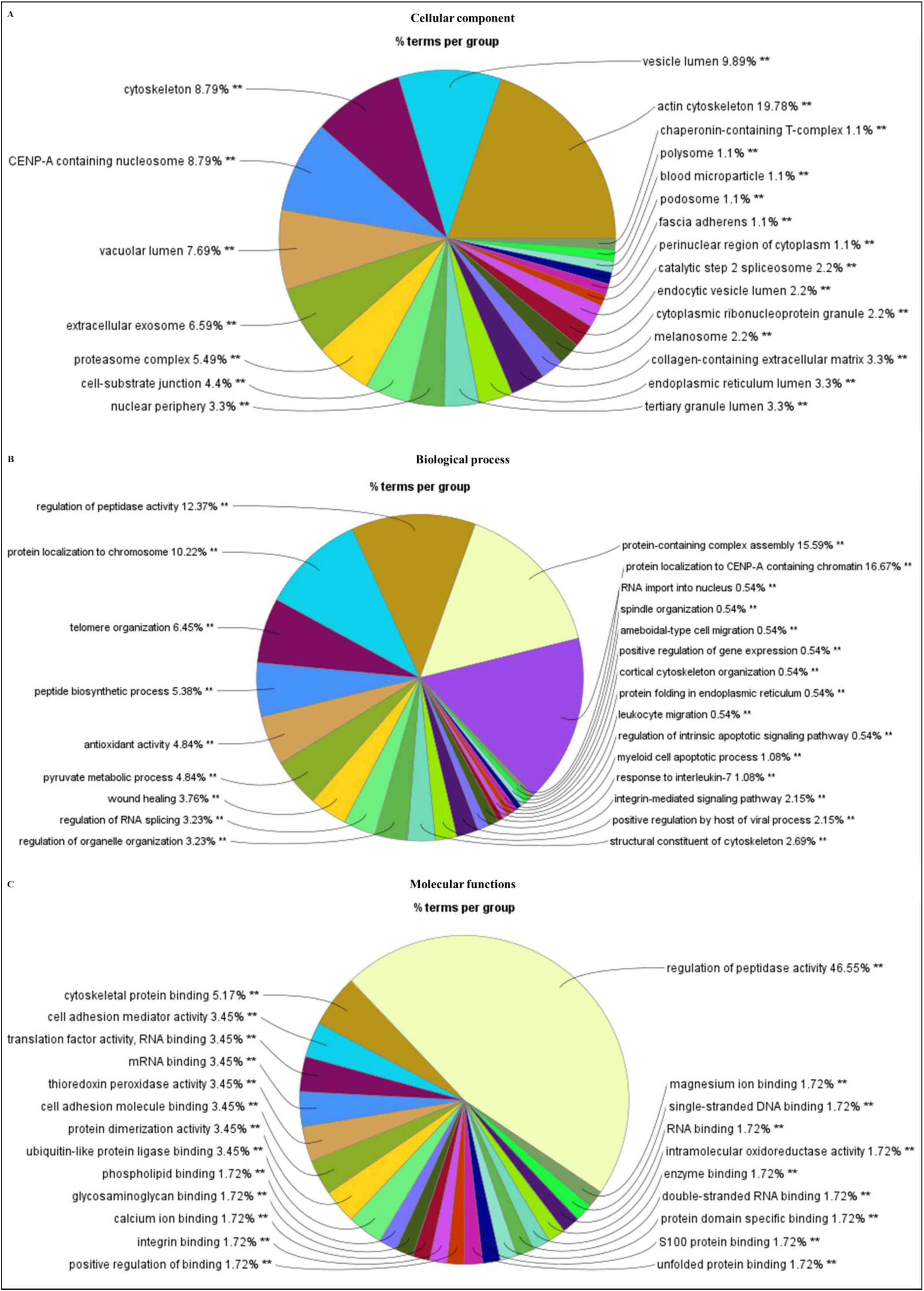
Gene Ontology (GO) analysis of upregulated proteins in co-culture of BM-MSCs and aPBMNCs. Pie charts depict (A) Cellular component. (B) Biological process. (C) Molecular function. Data showed a single independent experiment conducted with a single donor. **≤0.01

**Figure S5 (III):**
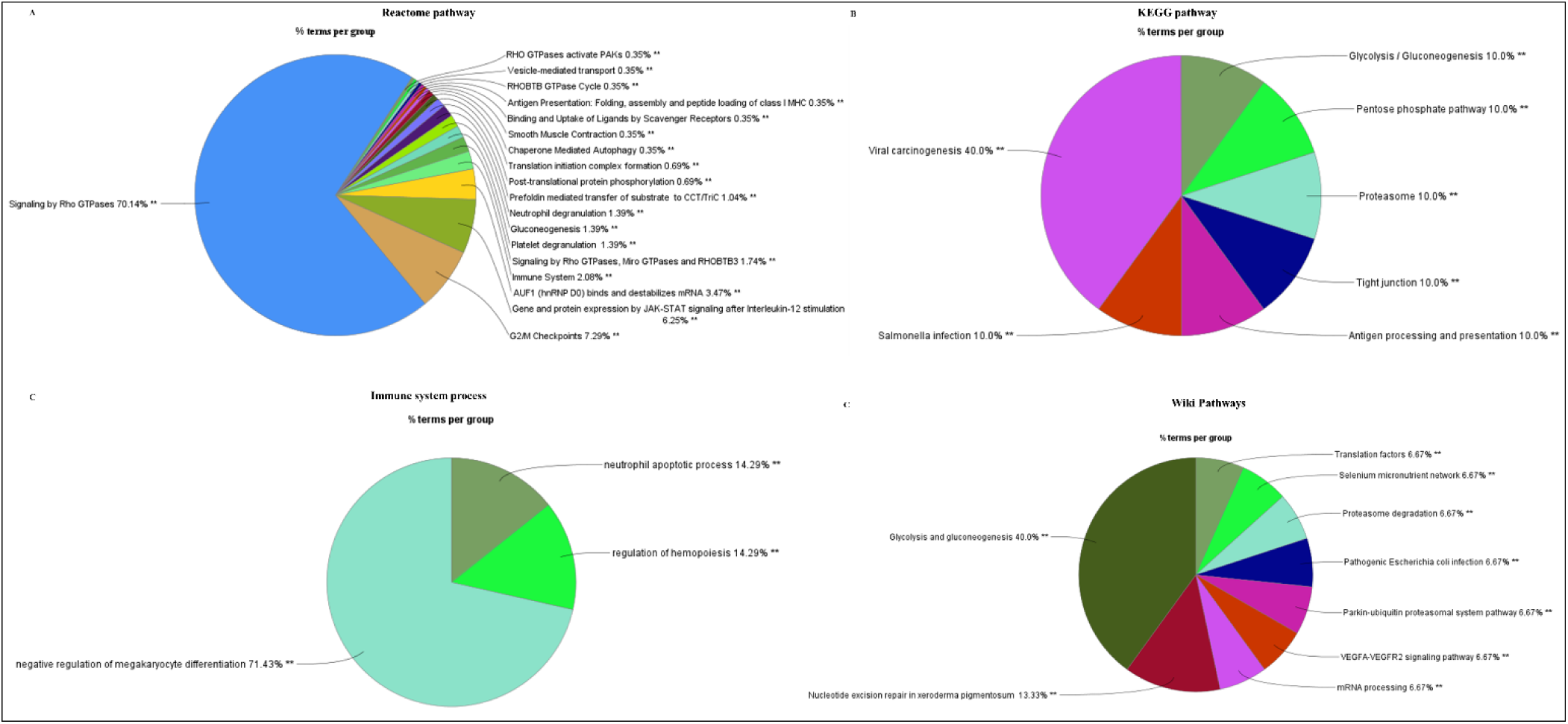
Functional enrichment analysis of upregulated proteins in co-culture of BM-MSCs and aPBMNCs. Pie charts depict (A) Reactome pathway (B) KEGG pathway (C) Immune system process. (D) WiKi pathway. Data showed a single independent experiment conducted with a single donor. **≤0.01

